# Meta-analyses of mouse and human prostate single-cell transcriptomes reveal widespread epithelial plasticity in tissue regression, regeneration, and cancer

**DOI:** 10.1101/2024.01.30.578066

**Authors:** Luis Aparicio, Laura Crowley, John R. Christin, Caroline J. Laplaca, Hanina Hibshoosh, Raul Rabadan, Michael M. Shen

## Abstract

Recent advances in single-cell RNA-sequencing (scRNA-seq) technology have facilitated studies of cell states and plasticity in tissue maintenance and cancer, including in the prostate. Here we present meta-analyses of multiple new and published scRNA-seq datasets to establish reference cell type classifications for the normal mouse and human prostate. Our analyses demonstrate transcriptomic similarities between epithelial cell states in the normal prostate, in the regressed prostate after androgen-deprivation, and in primary prostate tumors. During regression in the mouse prostate, all epithelial cells shift their expression profiles towards a proximal periurethral (PrU) state, demonstrating an androgen-dependent plasticity that is restored to normal during androgen restoration and regeneration. In the human prostate, we find progressive rewiring of transcriptional programs across epithelial cell types in benign prostate hyperplasia and treatment-naïve prostate cancer. Notably, we detect copy number variants predominantly within Luminal Acinar cells in prostate tumors, suggesting a bias in their cell type of origin, as well as a larger field of transcriptomic alterations in non-tumor cells. Finally, we observe that Luminal Acinar tumor cells in treatment-naïve prostate cancer display heterogeneous androgen receptor (AR) signaling activity, including a split between high-AR and low-AR profiles with similarity to PrU-like states. Taken together, our analyses of cellular heterogeneity and plasticity provide important translational insights into the origin and treatment response of prostate cancer.

## Introduction

Despite decades of investigation, regression and regeneration of the prostate gland as well as its oncogenic transformation represent fundamental biological processes that are poorly understood. In particular, androgen signaling represents a key regulatory program that maintains the identity of prostate tissue, yet the roles for androgen regulation in specific cell types remain unclear. In this regard, the advent of scRNA-seq technology has provided new tools to investigate the dynamics of prostate cell identity at the molecular level in both homeostasis and disease.

Although the prostate surrounds the urethra directly underneath the bladder, there are substantial anatomic differences between mammalian species. The mouse prostate is comprised of four distinct lobes, corresponding to the ventral (VP), lateral (LP), dorsal (DP), and anterior prostate (AP) lobes, whereas the human prostate lacks distinct lobular organization but can be subdivided into central, transition, and peripheral zones ***(Abate-Shen and Shen, 2000)***. These prominent anatomic differences have led in part to long-standing questions about the relationship of cell types and molecular pathways between the mouse and human prostate.

Classically, histological and ultrastructural analyses have described three major epithelial cell types in the prostate: luminal cells, basal cells, and rare neuroendocrine cells ***(Crowley and Shen, 2022; Toivanen and Shen, 2017)***, with less well-defined stromal cell types. However, recent scRNA-seq analyses have revealed considerable cellular heterogeneity and novel cell types in the mouse prostate epithelium ***(Crowley et al., 2020; Guo et al., 2020; Joseph et al., 2020; Karthaus et al., 2020; Mevel et al., 2020)***, and stroma ***(Joseph et al., 2021; Kwon et al., 2019)***. Although these studies independently reported multiple cellular populations with similar features, there are notable discrepancies in their nomenclature and description ***(Crowley and Shen, 2022)***, perhaps due to methodological differences in sample collection, preparation, computational analyses, and/or annotations. Similar issues also apply for scRNA-seq analyses focused on normal human prostate ***(Crowley et al., 2020; Guo et al., 2020; Henry et al., 2018; Karthaus et al., 2020)***, as well as in the context of pan-tissue resources ***(Eraslan et al., 2022; Tabula Sapiens et al., 2022)***. As a consequence, published scRNA-seq analyses of the mouse and human prostate are not readily comparable, and the precise relationships between cell populations described in different studies are unclear.

To address these issues, we have performed a meta-analysis of independent scRNA-seq datasets from the mouse prostate, aggregating datasets from two different mouse strains published by seven different laboratories and using two distinct bioinformatic approaches for their analysis to generate a comprehensive reference atlas. We have included new datasets to supplement rare cell populations, including a dataset of the proximal prostate (closest to the urethra) to examine the periurethral (PrU) cells residing in this region, and report gene signatures for each well-documented population during homeostasis. We have also analyzed time courses of prostate regression and regeneration, which demonstrate that each epithelial cell type displays similar transcriptomic shifts towards a PrU-like state following castration and returns to normal when androgen is reintroduced, revealing substantial androgen-dependent plasticity.

Similarly, we have performed a meta-analysis of the normal human prostate ***(Crowley et al., 2020; Guo et al., 2020; Joseph et al., 2020; Tabula Sapiens et al., 2022)*** to generate a consensus atlas of human prostate cell types during homeostasis. Since these studies have used different naming schemes and definitions for cell types, we have generated a nomenclature comparison and proposed a common descriptive naming scheme. In addition, we have compared the transcriptomic profiles of normal human and mouse epithelial cell types, and show that PrU cells in the human prostate have transcriptomic profiles consistent with reduced androgen sensitivity.

Finally, we have investigated changes in cell states that occur during progression to prostate adenocarcinoma. We have analyzed the dynamic changes in profiles of each cell type in homeostasis, hyperplasia ***(Joseph et al., 2020)***, and adenocarcinoma ***(Chen et al., 2021; Ge et al., 2022; Hirz et al., 2023; Karthaus et al., 2020; Song et al., 2022)***. Notably, we have found that Luminal Acinar (LumAcinar) cells display the greatest transcriptomic changes during progression to adenocarcinoma. Moreover, we found that copy number variants (CNVs) are only present in LumAcinar and rare neuroendocrine (NE) tumor cells, suggesting a predominant cell of origin for prostate cancer (PCa). However, we find that many LumAcinar cells lacking CNVs as well as other epithelial cell types also display extensive transcriptomic alterations, which is suggestive of a field effect similar to those observed in other tumor types. Most interestingly, we observe that tumor cells found in some treatment-naïve adenocarcinomas display a transcriptomic shift toward a PrU-like or LumDuctal state that displays decreased AR signaling activity. Taken together, our single-cell analyses demonstrate the cross-species conservation of prostate cell types, and underscore the significance of cellular plasticity following androgen deprivation as well as oncogenic transformation.

## Results

### Epithelial populations of the mouse prostate

To generate a comprehensive aggregated single-cell dataset for the mouse prostate, we gathered publicly available scRNA-seq datasets generated from C57BL/6 and FVB mice, and also generated new data focusing on the proximal and periurethral regions of the prostate, which have been less studied. We analyzed these datasets using two independent computational approaches to confirm the reproducibility of our interpretations. In the first approach, we de-noised each dataset using Random Matrix Theory (RMT), which improves the ability to separate and detect rare cell populations ***(Aparicio et al., 2020)***. We then sequentially clustered in each dataset to identify cell populations, following the same strategy used previously ***(Crowley et al., 2020)***. This analysis of 24 datasets resulted in an aggregated dataset of 135,831 cells arranged in 30 clusters (Figure 1A; Figure 1—figure supplements 1A, 2). As a second approach, we used a standard Seurat pipeline to generate an aggregated dataset from 13 datasets of sufficiently high quality, which was composed of 30,433 cells in 18 distinct prostate cell clusters (Figure 1–figure supplement 3A).

**Figure 1.**
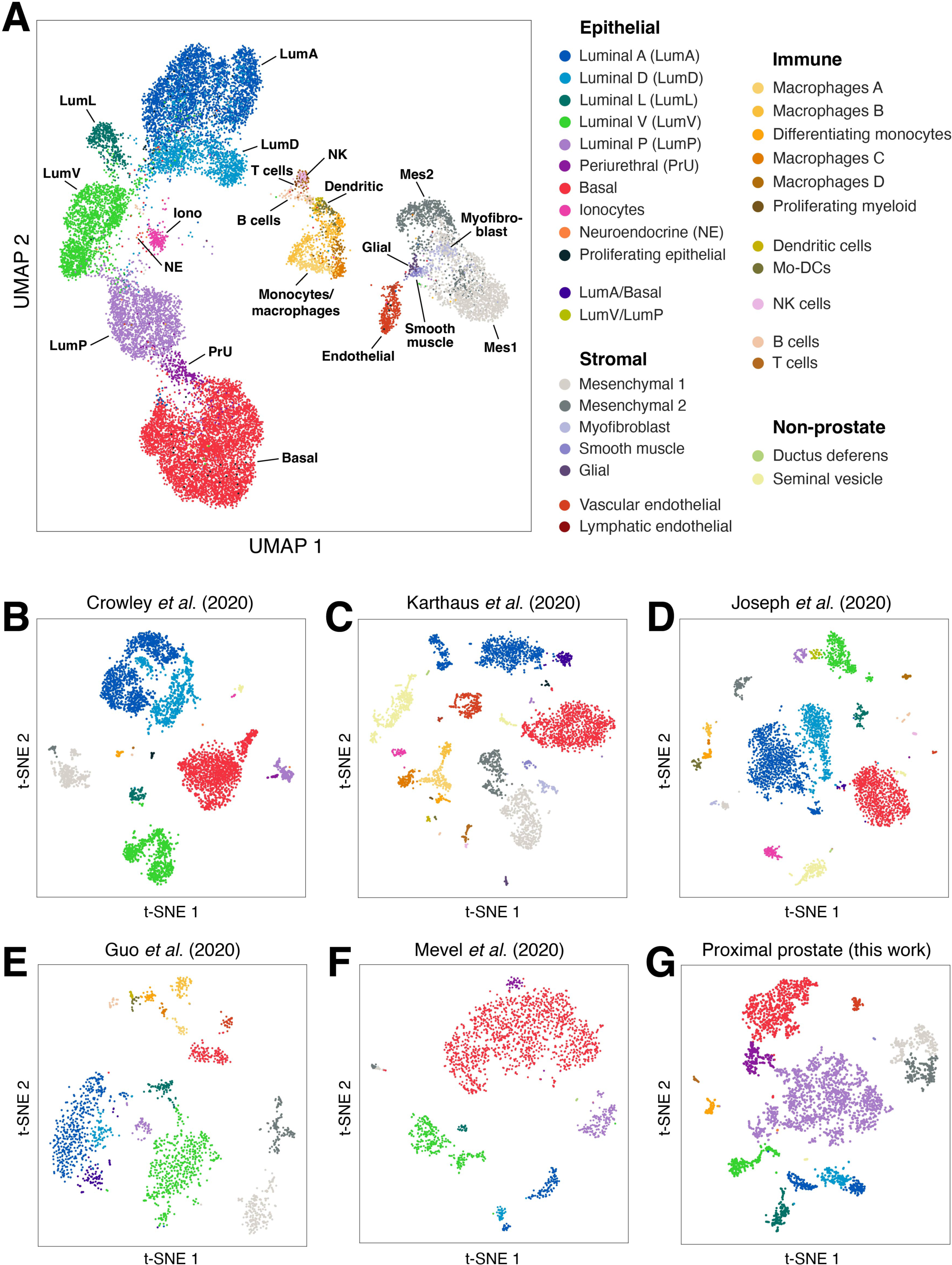
Reference plots of mouse prostate scRNA-seq data demonstrate extensive cell type heterogeneity. Shown are an aggregated composite plot **(A)**, as well as plots of individual datasets corresponding to: C57BL/6 whole prostate ***(Crowley et al., 2020)* (B)**, FVB anterior prostate lobe ***(Karthaus et al., 2020)* (C)**, C57BL/6 whole prostate ***(Joseph et al., 2020)* (D)**, C57BL/6 whole prostate ***(Guo et al., 2020)* (E)**, C57BL/6 whole prostate ***(Mevel et al., 2020)* (F)**, and C57BL/6 proximal prostate (this work; **G**). Datasets were processed using the *Randomly* pipeline, which revealed 12 epithelial populations, 7 stromal populations, and 11 immune populations found across multiple datasets. Non-prostatic populations as well as populations that may correspond to cell states are only shown for the individual datasets.

These parallel approaches allowed for the identification and comparison of cell populations across datasets in a uniform manner, independent of differences in reporting and labeling between publications. To define robust cell populations, we required that the population be identified in at least three independent datasets and have nearly complete overlap in globally distinguishing gene expression. For rare cell populations, we only required that the population be present in at least two independent datasets. Of note, the majority of clusters were identical using both the RMT and Seurat approaches. The RMT approach handled sparse data differently, yielding a greater number of small clusters and providing better discrimination between populations with low cell numbers.

We found that epithelial populations were remarkably consistent across datasets and approaches. Interestingly, no distinct subclusters were formed based on mouse strain background, which did not significantly contribute to prostate epithelial heterogeneity. In particular, basal cells formed a single contiguous cluster in individual datasets (Figure 1B-F; Figure 1—figure supplement 1B-F), as previously reported ***(Crowley et al., 2020; Guo et al., 2020; Joseph et al., 2020; Karthaus et al., 2020; Mevel et al., 2020)***, and in our aggregated datasets (Figure 1A; Figure 1—figure supplement 1A). We did not observe evidence of a distinct basal subcluster with expression of Zeb1 or other epithelial-mesenchymal transition (EMT) markers ***(Wang et al., 2020)***. However, in the Seurat pipeline, we observed that a small subset of basal cells adjoins the periurethral (PrU) cluster and expresses slightly more proximal and luminal markers (Figure 1—figure supplement 1A).

We identified multiple luminal epithelial clusters, which represent distinct cell types that are separated by prostate region (Figure 1A-F; Figure 1—figure supplement 1A-F), as previously reported for individual datasets ***(Crowley et al., 2020; Guo et al., 2020; Joseph et al., 2020; Karthaus et al., 2020; Mevel et al., 2020)***. Since the nomenclature for these populations differs between laboratories (summarized in ***(Crowley and Shen, 2022)***), we follow a descriptive naming system ***(Crowley et al., 2020)*** that denotes lobe-specific prostate populations (*e.g.,* LumA for the distal anterior lobe) as well as proximal populations (LumP for proximal prostate). Notably, although the dorsal and lateral lobes have often been combined as a “dorsolateral lobe”, highly distinct dorsal (LumD) and lateral (LumL) populations were always found in each individual dataset as well as the aggregated datasets (Figure 1A,B,D-F; Figure 1—figure supplement 1A,B,D-F). In contrast, the anterior (LumA) and dorsal (LumD) distal luminal populations consistently displayed the most transcriptomic overlap (Figure 1A,B,D-F; Figure 1—figure supplement 1A,B,D-F).

Unlike distal luminal cells, which differ by lobe, proximal luminal cells (LumP) formed a single cluster without lobe-specific identity (Figure 1A-F; Figure 1—figure supplement 1A-F) ***(Crowley et al., 2020; Guo et al., 2020; Joseph et al., 2020; Karthaus et al., 2020; Mevel et al., 2020)***. The vast majority of LumP cells are located in the proximal region of the prostate, though rare distal cells can be observed ***(Crowley et al., 2020; Guo et al., 2020; Mevel et al., 2020)***, and functional heterogeneity within the population has been reported ***(Guo et al., 2020)***. In this regard, in the Seurat pipeline, we observed that a subset of LumP cells clustered closer to distal luminal cells (Figure 1—figure supplement 1A).

Neuroendocrine (NE) cells represent a rare and historically elusive epithelial population that could be detected in both analytical pipelines (Figure 1, Figure 1—figure supplement 1). Interestingly, NE cells can co-express distal luminal (LumDist), basal, LumP, or PrU markers, suggesting population heterogeneity ***(Crowley et al., 2020; Guo et al., 2020; Joseph et al., 2020)***. Ionocytes are another rare population that was recently described in the prostate ***(Karthaus et al., 2020)***, and our meta-analysis revealed their presence in additional datasets (Figure 1A,B,D; Figure 1—figure supplement 1A,B,D) ***(Crowley et al., 2020; Joseph et al., 2020)***. Though ionocytes have some transcriptional similarities to NE cells, they express *Foxi1* and *Atp6v1g3* but not specific luminal or basal markers (Figure 1—figure supplement 1I,J). Both cell types were observed in higher proportions in the proximal dataset, and the PrU population is described in detail below.

Using our aggregated datasets, we generated reference gene expression signatures that are specific for each prostate epithelial cell type (Supplementary file 1). In addition, we examined the Gene Set Enrichment Analysis Hallmark signatures and found increased expression of genes involved in protein secretion in seminal vesicle and distal luminal cells, and the lowest levels of Notch signaling genes in NE cells (Figure 1––figure supplement 4). Finally, we observed rare epithelial clusters in individual datasets that may represent cell states. In particular, a subset of LumA cells expresses both LumA and Basal markers, and may correspond to “intermediate” cells with hybrid luminal and basal features (Figure 1—figure supplement 1B,C).

### Non-epithelial cell populations

Our scRNA-seq meta-analysis also provided consistent insights into non-epithelial cell types in the mouse prostate. The mesenchymal/stromal cells present in these datasets are predominantly fibroblasts, and can be divided into several different clusters (Figure 1; Figure 1—figure supplements 2E,F, 3). The Mesenchyme 1 (Mes1) population is proximally-enriched, lies adjacent to the epithelium, and expresses *Srd5a2* as well as many Wnts and other signaling factors, whereas Mesenchyme 2 (Mes2) is enriched more distally, is located slightly farther from the epithelium, and expresses many chemokines and complement components ***(Crowley et al., 2020; Kwon et al., 2019)***. We also identified distinct myofibroblast and smooth muscle populations that express smooth muscle actin (*Acta2*), and observed that a subset of myofibroblasts expresses *Lgr5 **(Crowley et al., 2020; Karthaus et al., 2020; Kwon et al., 2019; Wei et al., 2022)***. Although a third fibroblast population has been reported (Kwon et al., 2019), it did not appear as a distinct cell type in our analyses, but rather as a subset of Mes2 (Figure 1—figure supplement 1E,F). Interestingly, several mesenchymal cell types reported to exist in the prostate (*e.g.,* telocytes) were not detected in any dataset, suggesting that the prostate stromal compartment is incompletely captured in existing scRNA-seq data.

Hematopoietic lineage populations (such as B and T lymphocytes, dendritic cells, and NK cells) were also detected across multiple datasets, with the immune compartment displaying a notable myeloid bias. In particular, macrophages divided into distinct subclusters along a continuous spectrum, which was most evident in the RMT pipeline. Since profiles for M1 and M2 macrophages could not be definitively identified, we have named these populations alphabetically (Figure 1; Figure 1—figure supplement 1A-D). In addition to the macrophage populations, we detected a population with substantial overlap in gene expression to macrophages, which appeared to correspond to differentiating monocytes (Figure 1—figure supplement 1A-D).

Finally, we also observed contaminating seminal vesicle cells across multiple datasets. Seminal vesicle epithelial cells could be clustered into a single basal population as well as luminal populations with more proximal markers or more distal markers (Figure 1—figure supplement 1G,H), suggesting potential epithelial heterogeneity within this tissue.

### The periurethral region

We define the periurethral (PrU) region as the most proximal extent of each prostate lobe nearest the junction with the urethra. PrU cells make up most of the epithelium in this region. Because this region is located exclusively within the rhabdosphincter and hence is more difficult to dissect, many prostate scRNA-seq samples have not captured the epithelial populations in this region. However, our meta-analysis detected PrU epithelial cells in several datasets ***(Crowley et al., 2020; Joseph et al., 2020)*** as well as many in our proximal prostate scRNA-seq dataset (Figure 1G; Figure 1—figure supplement 1G). Uniquely, PrU epithelial cells display hybrid luminal and basal features, similar to urothelial cells in the adjacent urethra. However, PrU cells can be readily distinguished from urethral cells by lineage-tracing with an *Nkx3.1-Cre* driver ***(Crowley et al., 2020)***.

To understand the unique morphological features of PrU cells, we imaged the periurethral region by electron microscopy and immunofluorescence staining (Figure 2). At the ultrastructural level, PrU cells share some features with distal luminal (LumDist) cells, such as organelles involved in protein secretion, and many features with LumP cells, including a high density of mitochondria (Figure 2A-F). Interestingly, several features of PrU cells also resemble urothelial cells of the urethra, including the nuclear orientation of more basally-situated PrU cells, as well as the lumen-facing structures of apically-situated cells, which may resemble the rigid, uroplakin-filled surface of urothelial cells. Thus, PrU cells share ultrastructural features of both the prostate and the urethral urothelium, and may represent a physical transition between the two tissues.

**Figure 2.**
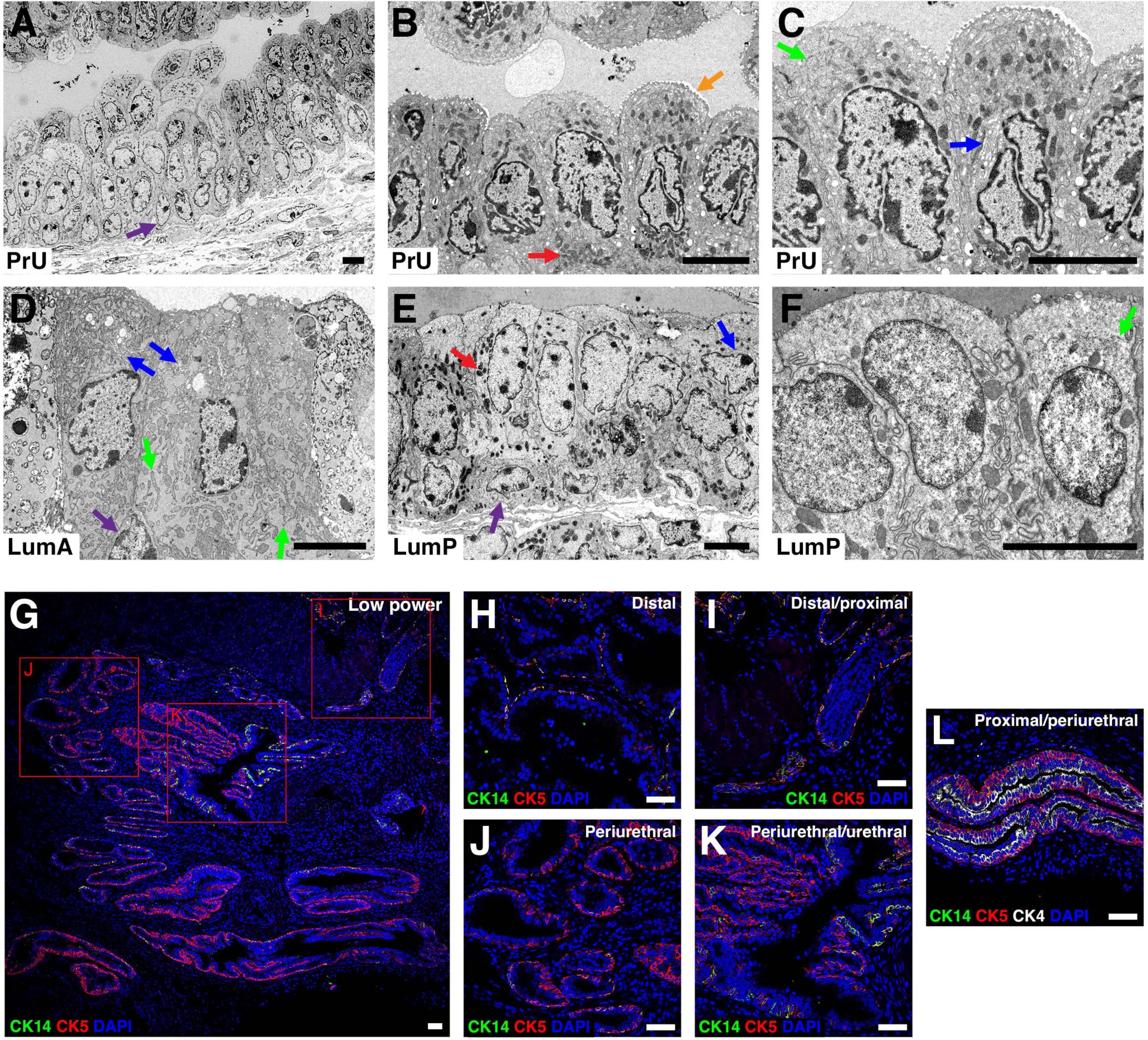
Imaging of mouse PrU cells reveals unique and shared features with prostatic and urethral cells. **(A-C)** Scanning electron microscopy (EM) images of PrU cells show a focal region of cells where they appear to be multilayered **(A)**, a region that is not multilayered and displays unique features **(B)**, and a higher magnification of this region **(C)**. **(D-F)** The features of distal LumA cells **(D)**, proximal LumP and basal cells **(E)**, and LumP cells at higher magnification **(F)** are shown for comparison. Arrows indicate basal nuclear orientation (purple), mitochondrial density (red), apical membrane structures (orange), rough endoplasmic reticulum (green), and Golgi apparatus (blue). Scale bars in **A-F** indicate 5 microns. **(G-L)** Immunofluorescence staining show changes in basal and proximal keratin expression. (G) Overview of the periurethral region with neighboring urethral and proximal cells at low power. Insets show co-expression of basal keratins CK5 (red) and CK14 (green) in distal **(H)** and proximal **(I)** basal cells, and consistent CK5 but reduced CK14 in periurethral **(J)** and periurethral and urethral **(K)** basal cells. Proximal keratin CK4 (white) is maintained through the proximal and periurethral region **(L)**. No superficial-like cells were observed in the periurethral region. Scale bars in **G-L** indicate 50 microns.

At the level of gene expression, PrU cells uniquely express *Lmo1, Anxa8, Dapl1,* and *Aqp3,* and have higher *Ly6d* and *Sca-1* expression than LumP cells ***(Crowley et al., 2020)*** (Figure 2—figure supplement 2B). Although *Krt5* and *Krt14* expression overlaps in the basal layer throughout more distal regions of the prostate, *Krt14* expression becomes intermittent in the PrU region and *Krt5* is maintained, whereas basal cells of the urothelium rarely express *Krt14* (Figure 2G-L). Based on our re-analysis of a scRNA-seq dataset of the proximal prostate and urethra ***(Joseph et al., 2020)***, we could define two distinct urethral populations, a luminal-intermediate urothelial cell group with transcriptomic similarity with LumP cells, and a basal-intermediate urothelial cell group with similarity with PrU cells (Figure 2—figure supplement 2). Notably, at homeostasis, PrU and LumP cells can be readily distinguished from urothelial cells by key markers, including several uroplakins (Figure 2—figure supplement 2). Thus, PrU cells also represent a transition population in terms of molecular features, such as gene expression.

### The transcriptomic response to androgen deprivation and restoration

The prostate regresses in response to androgen-deprivation and regenerates after androgen restoration, which can be repeated through at least 30 cycles in the mouse ***(Isaacs, 1985; Tsujimura et al., 2002)***. To examine the response of individual cell populations to androgen-deprivation and restoration, we examined scRNA-seq data of mouse prostate through time courses of regression and regeneration ***(Guo et al., 2020; Karthaus et al., 2020)***. For this analysis, we defined a “cell type score” to represent the average of the most specific and differentially expressed genes for each cell type (Methods). In response to castration, every cell type except endothelial cells showed a significant decrease in its cell type score (Figure 3A,B). Interestingly, the rates of transcriptomic change were different for each population, as distal luminal (LumDist) cells, myofibroblasts, and Mes1 cells rapidly lost almost all of their cell-type specific gene expression, whereas LumP, basal, smooth muscle, and Mes2 cells only lost approximately half of their cell-type specific gene expression, with Mes2 cells retaining their gene expression profile the longest (Figure 3A,B).

**Figure 3.**
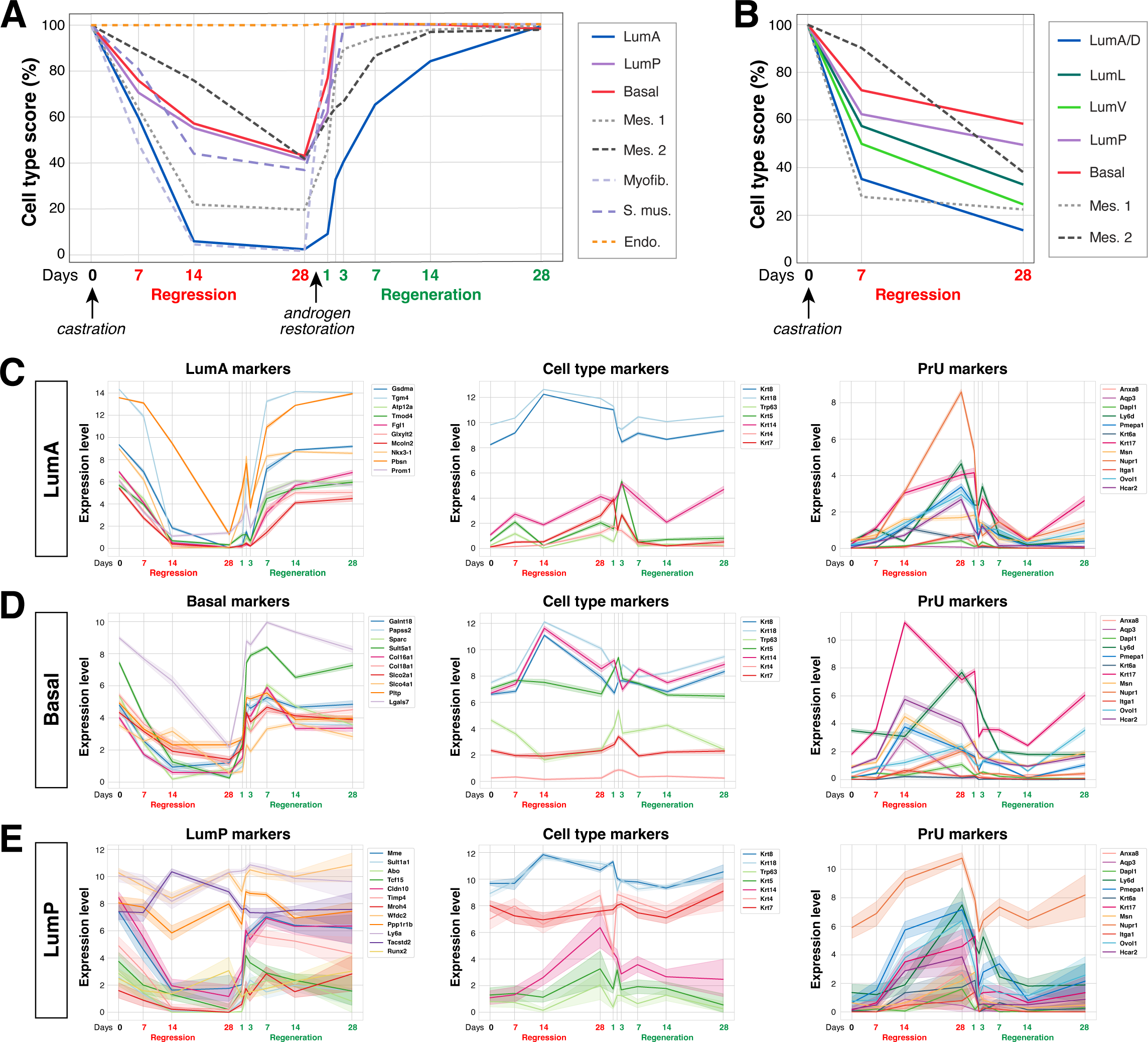
Time course of prostate regression and regeneration reveals androgen-dependent plasticity. **(A,B)** Meta-analyses of single-cell RNA-seq datasets for prostate regression and regeneration based on ***(Karthaus et al., 2020)*** (A) and for regression based on ***(Guo et al., 2020)*** (B). “Cell type score” is defined as the percentage of most specific differentially expressed genes for each population, averaged over the whole population (Methods). **(C-E)** Changes in gene expression that are enriched in urethral but not PrU cells, such as *Areg* and *Ociad2*, in the LumA **(C)**, Basal **(D)**, and LumP **(E)** populations, showing distinguishing genes for each population (left column), genes for general compartmental markers, and genes that are enriched for PrU and not co-expressed in LumP (right column), where the line indicates the average expression for each gene across the population and the bar indicates confidence interval (+/- 95%).

Interestingly, our analysis indicated that mouse prostate epithelial cells shift toward a PrU-like expression profile during regression. A detailed examination of gene expression patterns in LumA, Basal, and LumP populations showed that each population lost expression of many specific genes but retained its distinctive expression of select distal luminal, basal, or proximal luminal markers during the regression-regeneration cycle (Figure 3C-E). However, each epithelial population gained expression of multiple PrU markers following castration and lost this expression after androgen restoration; furthermore, the markers retained by LumP cells during regression were those that are co-expressed by PrU cells. The epithelial populations did not shift towards urethral gene expression profiles, as only rare LumP cells expressed any urothelial markers (Figure 3—figure 3 supplements 1, 2). Notably, while the normal PrU profile includes some genes that are co-expressed by either LumP or the urethral urothelium, the regressed epithelium expresses many PrU-specific genes that are distinct from both. These findings highlight PrU-like transcriptomic profiles and provide a broader context for the previously reported shift from LumA towards LumP in the anterior prostate following androgen deprivation ***(Karthaus et al., 2020)***.

A transcriptomic shift was also observed in the prostate stroma during regression, as both the Mes1 and Mes2 fibroblast populations altered gene expression in response to androgen deprivation. Mes1 cells rapidly shifted toward a Mes2 expression profile and lost expression of several defining factors including Wnts, whereas Mes2 cells changed gene expression more slowly (Figure 3A,B; Figure 3—figure supplements 1, 2). Thus, we conclude that transcriptomic reprograming following androgen deprivation is not exclusive to the luminal or distal compartments, but instead represents a tissue-wide alteration of cell states.

### Atlas of the human prostate

Next, we performed a meta-analysis of published scRNA-seq datasets to establish a corresponding reference atlas of the normal human prostate ***(Crowley et al., 2020; Guo et al., 2020; Henry et al., 2018; Tabula Sapiens et al., 2022)*** using the criteria described for the RMT pipeline (Methods). Despite differences in the relative proportions of cell populations between these datasets, the data were remarkably consistent. We found that the human prostate has a single Basal epithelial population, two luminal populations corresponding to Luminal Acinar (LumAcinar) and Luminal Ductal (LumDuctal), and a Periurethral-like (PrU) population (Figure 4A-F; Figure 4—figure supplement 4A). The stromal populations were more variable and less well-represented across datasets, but corresponded to at least 1 endothelial population and 3 fibroblast-like populations (Figure 4A; Figure 4—figure supplement 4A,B). Of the 3 fibroblast-like populations, the first expressed several classic fibroblast markers and did not subdivide readily (we denote these as general fibroblasts), the second corresponded to fibroblasts that express several muscle genes (myofibroblasts), and the third to fibroblast-like cells that express many contractile muscle genes (fibromyocytes) (Figure 4F) ***(Travaglini et al., 2020)***. Based on differential gene expression, we generated signatures for each epithelial and mesenchymal population (Supplementary file 2). Within the immune compartment, we detected relatively fewer cells with variable representation of cell types between patients, so these populations were grouped as either myeloid or lymphoid. Interestingly, the zone of the prostate tissue did not have a clear effect on the transcriptome (Figure 4—figure supplement 4B,C), as previously reported ***(Guo et al., 2020)***.

**Figure 4.**
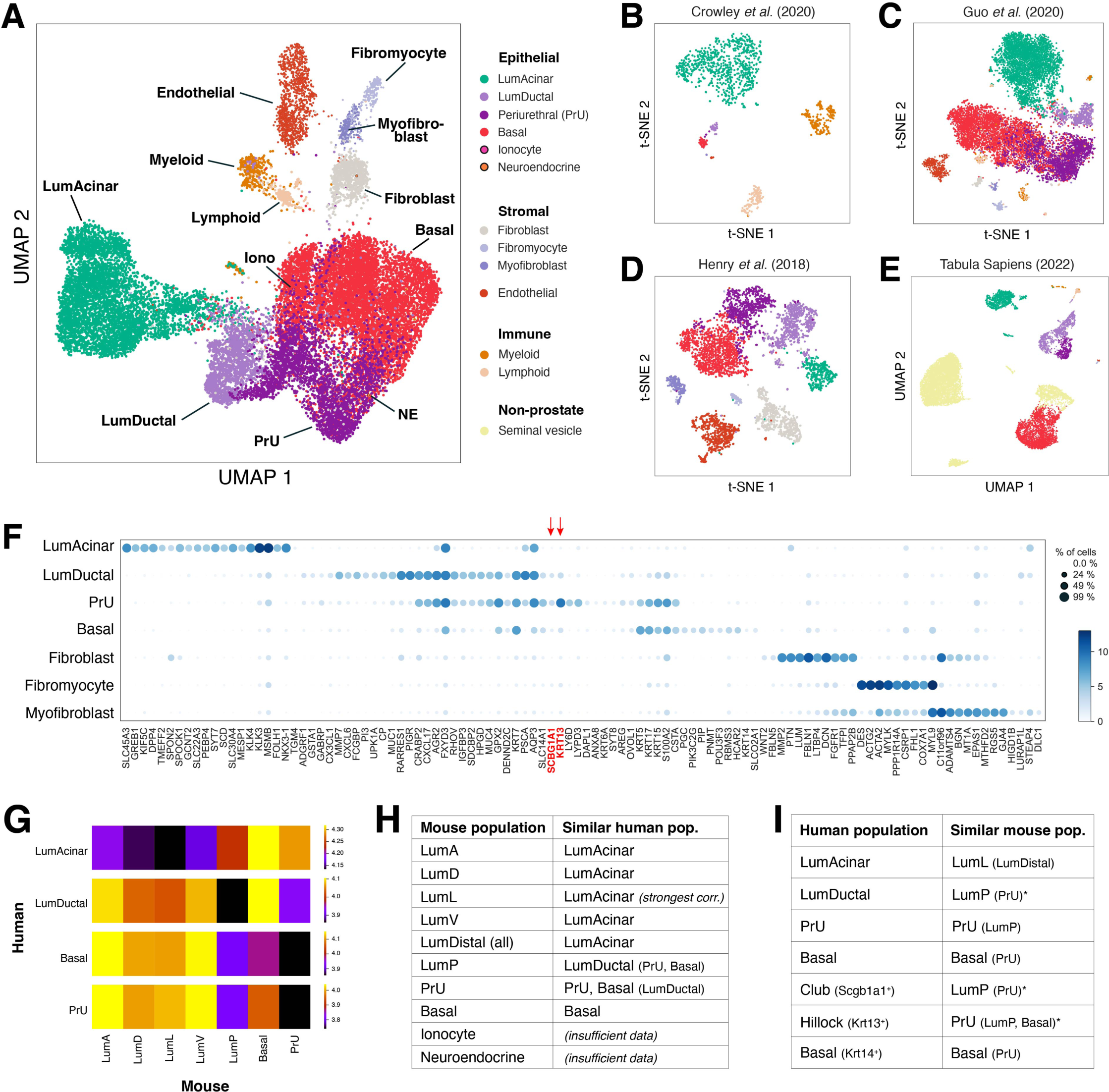
Reference plots for human prostate scRNA-seq data. Shown are an aggregated composite plot **(A)**, as well as the plots of individual datasets **(B-E)** for samples of benign human prostate and adjacent benign prostate. **(B)** UMAP plot corresponding to primarily LumAcinar cells taken from the peripheral zone of 1 patient ***(Crowley et al., 2020)***. **(C)** Plot containing primarily Basal and PrU cells from 1 patient with PCa ***(Guo et al., 2020)***. **(D)** Dataset containing primarily Basal, PrU, and LumDuctal cells from 1 patient without PCa ***(Henry et al., 2018)***. **(E)** Dataset containing mixture of prostate and seminal vesicle, originating from 2 organ donor patients with no history of prostate disease ***(Tabula Sapiens et al., 2022)***. **(F)** Dot plot of differentially expressed genes for the epithelial and stromal cell populations from the reference aggregated normal human prostate. The club cell marker SCGBA1 and hillock cell marker KRT13 are highlighted. **(G)** Heatmap comparing the total gene expression profiles of the cell types in the normal human prostate dataset ***(Tabula Sapiens et al., 2022)*** to those of the aggregated normal mouse prostate, using Wasserstein distance as a metric. Darker color indicates greater transcriptomic similarity. **(H, I)** Tables listing the most similar mouse and human epithelial populations based on gene expression, generated by overlaying the mouse cell type signatures onto the human populations **(H)** and vice versa **(I)**.

Since the nomenclature of human prostate epithelial populations differs between publications, we compared our previous nomenclature ***(Crowley et al., 2020)*** to an alternative system that uses “Club” and “Hillock” lung terminology ***(Henry et al., 2018)***, using the Tabula Sapiens as a source of normal tissue (Figure 4E; Figure 4—figure supplement 4D,E). Notably, we found that most of the “Club” cells corresponded to LumDuctal and PrU cells (Figure 4—figure supplement 4B,C), as they expressed common luminal genes and more specific markers like *RARRES1*, but did not consistently express the defining marker *SCGB1A1* (Figure 4F). Similarly, most “Hillock” cells corresponded to PrU cells (Figure 4—figure supplement 4B,C), as they expressed common luminal and basal genes as well as more specific markers such as *KRT7*, *PSCA*, *RARRES1*, *LYPD3*, and *AQP3*; moreover, expression of the Club- and Hillock-defining markers were not specific (Figure 4). The remaining luminal cells corresponded to LumAcinar cells (Figure 4—figure supplement 4D,E), expressing common luminal cytokeratins as well as more specific markers including *KLK3*, *MSMB*, *FOLH1*, and *TGM4* (Figure 4F). These transcriptional similarities were separately confirmed by examination of single-nuclear RNA-seq data from the GTEx project ***(Eraslan et al., 2022)***. Based on these analyses, we find that our descriptive nomenclature of human prostate epithelial populations correlates with lung terminology but appear to align more accurately with distinct cell types in the prostate.

To perform an updated cross-species comparison of cell type identities ***(Crowley et al., 2020)***, we calculated the Wasserstein distance between gene expression profiles for each population in the aggregated mouse and human datasets in transcriptomic latent space (Methods) (Figure 4G). While human and mouse Basal cells have notably different profiles, human Basal and PrU populations most closely resemble mouse PrU, human LumDuctal most closely resembles mouse LumP, and human LumAcinar most closely resembles mouse LumL followed by LumD (Figure 4—figure supplement 4D). To test the robustness of this analysis, we removed individual genes from the mouse expression profiles and repeated the comparisons, which revealed that the greater similarity of human LumAcinar to LumL was dependent on expression of a handful of specific distinguishing genes, such as *Msmb* in LumL, and that the transcriptomes of the LumDist populations had mostly similar marker overlap with human LumAcinar otherwise. Consequently, we suggest that human LumAcinar cells, which are distributed throughout different zones of the human prostate, correspond more generally to mouse LumDist populations of all lobes. We additionally plotted the signatures of each population on the aggregated data of the other species to see how the differentiating genes versus the whole transcriptome compare across species; these results were summarized as tables (Figure 4H,I). Together, these results suggest a clear correlation across species.

### Distinguishing human prostate cancer progression by AR signaling levels

To examine the alterations of the human prostate due to disease, we combined the normal prostate scRNA-seq datasets with those from patients with benign prostate hyperplasia (BPH) ***(Joseph et al., 2020)***, and treatment-naïve prostate cancer ***(Chen et al., 2021; Ge et al., 2022; Hirz et al., 2023; Karthaus et al., 2020; Song et al., 2022)***. In these aggregated data of 99,611 cells from 66 datasets, we observed heterogeneous gene expression profiles across treatment-naïve tumors, which was particularly apparent in LumAcinar cells from prostatectomy samples. Therefore, we performed PHATE visualization of the LumAcinar cells from the aggregated data to depict local and global data structures (Figure 5A; Figure 5—figure supplement 5A,B). This analysis revealed that LumAcinar cells across early prostate disease stages can be subclustered into six primary groups according to disease stage with different gene expression profiles (Figure 5B). Notably, these groups divide into two distinct arms that correlate with androgen receptor (AR) signaling levels and PrU-like gene expression. Pseudotime analysis suggested that the AR-high and AR-low arms may both initiate from normal (from healthy prostates) and/or intermediate stages (normal-like gene expression from prostatectomy samples); furthermore, they may pass through a hyperplastic expression stage before splitting into distinct arms (Figure 5C).

**Figure 5.**
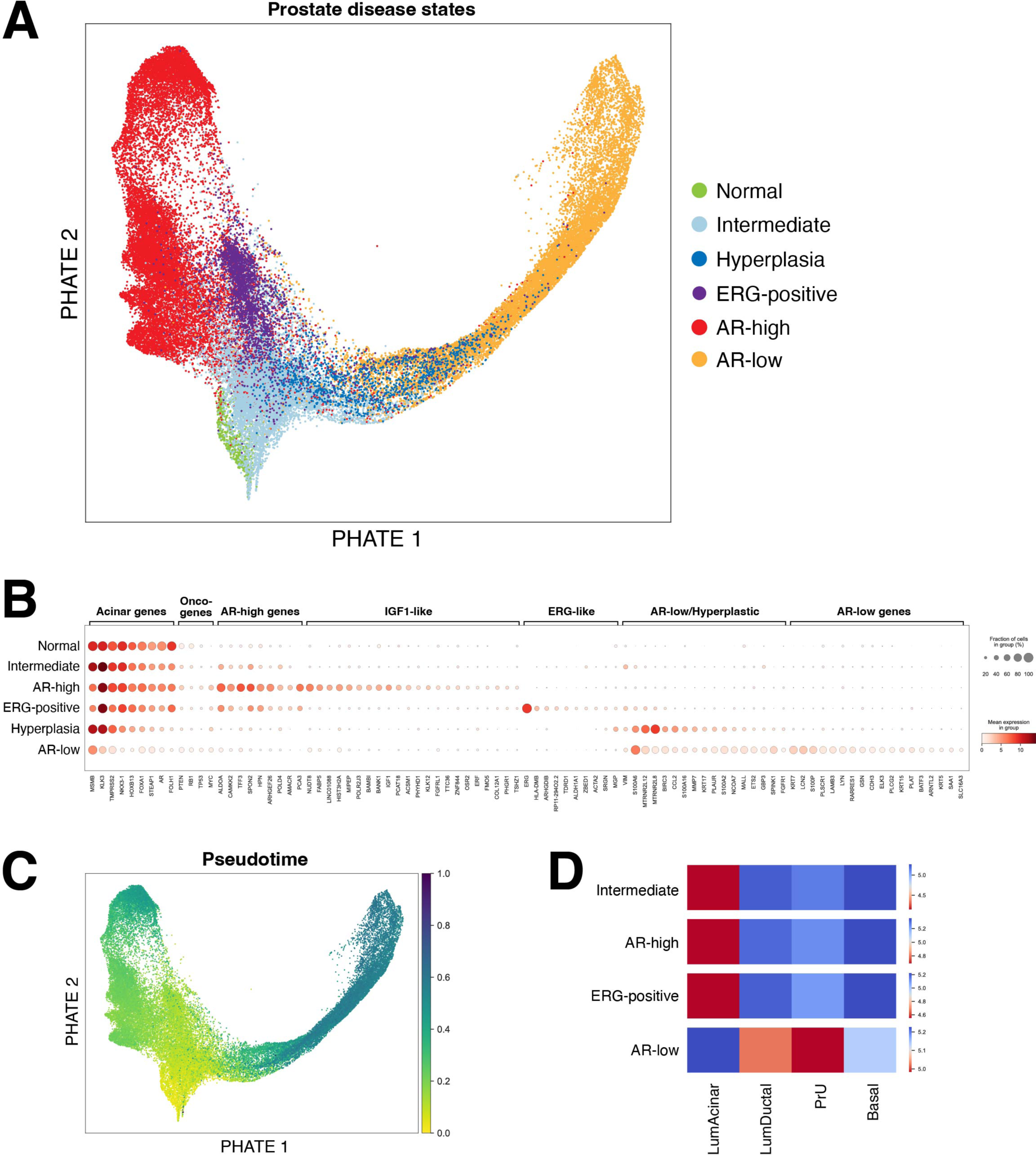
Meta-analysis of scRNA-seq datasets from human prostate adenocarcinoma reveals disease evolution of Luminal Acinar cells. **(A)** PHATE plot of LumAcinar populations from 2 normal prostates *(Tabula Sapiens et al., 2022)*, 3 prostates with BPH *(Joseph et al., 2020)*, and 46 prostates with PCa from Memorial Sloan Kettering Cancer Center *(Karthaus et al., 2020)*, University of California, San Francisco *(Song et al., 2022)*, Massachusetts General Hospital *(Hirz et al., 2023)*, Peking University Third Hospital *(Ge et al., 2022)*, and Shanghai Changhai Hospital *(Chen et al., 2021)*. Clustering of the aggregated data reveals that LumAcinar cells show the greatest variation, as LumAcinar cells from the normal and BPH prostates occupy 1 cluster each (Normal and Hyperplasia, respectively), while cells in the PCa samples subdivide into 4 major subpopulations (Intermediate, ERG-positive, AR-high, and AR-low). The PHATE plot splits these PCa subpopulations along two major branches. **(B)** Dot plot of cell type, PCa, subpopulation-defining, and other relevant genes indicates that AR signaling is a major differentiating factor across the two branches. **(C)** Pseudotime analysis of the cells in **A** suggests a Normal or Intermediate origin for both branches in PCa. This also suggests progression through Hyperplasia to PCa in some cells. **(D)** Heatmap comparing the total gene expression profiles of the LumAcinar subpopulations in **A** to all normal epithelial populations. The AR-low branch has a notable shift towards PrU and LumDuctal marker expression. Wasserstein distance is used as a metric, and darker red indicates greater transcriptomic similarity.

We performed several analyses to understand this division of gene expression profiles during prostate cancer progression. First, we compared the profiles of the four LumAcinar groups found in PCa samples to profiles of normal human prostate epithelial populations. We found that the cells of the AR-high and ERG-positive groups as well as the intermediate group resembled normal LumAcinar cells and retained the expression of many differentiated LumAcinar genes (Figure 5B,D). In contrast, LumAcinar cells of the AR-low group shifted from LumAcinar toward PrU expression patterns (Figure 5B,D). Additional genes that are lost or gained during the transition from normal LumAcinar to AR-high or AR-low tumor cells were also noted (Figure 5—figure supplement 5D). To confirm this analysis, we mapped a signature of the most differentially expressed genes in the AR-low arm, as well as the Hallmark AR response signature and an independent AR response signature ***(Spratt et al., 2019)*** onto the aggregated LumAcinar populations (Figure 5—figure supplement 5). Together, these data indicate that LumAcinar cells in primary treatment-naïve prostate cancer can be divided into two primary groups of gene expression patterns based on AR signaling levels. AR-high and ERG-positive cells display elevated AR signaling relative to normal LumAcinar cells and are associated with classical PCa features. In contrast, AR-low cells have dramatically reduced AR signaling levels and shift toward PrU and some LumDuctal expression profiles, unlike other transformed groups.

In addition, our analyses identified two rare and distinct subsets of LumAcinar cells that display markers of partial neuroendocrine differentiation (Figure 5—figure supplement 5A). One group expresses genes such as ASCL2 and POU2F3 and is located in the AR-low arm (Figure 5—figure supplement 5B). The other group expresses genes including ONECUT2 and INSM1, and is located predominantly within the ERG-positive subset in the AR-high arm (Figure 5—figure supplement 5C). Interestingly, these two groups may represent the early emergence of neuroendocrine transdifferentiation from luminal adenocarcinoma cells, corresponding to the Class 1 and 2 pathways, respectively, which were recently defined in analyses of a model of prostate neuroendocrine differentiation ***(Chen et al., 2023)***.

### CNVs are specific for LumAcinar cells

For robust identification of definitive tumor cells in the human prostate cancer scRNA-seq datasets, we used InferCNV to identify copy number variants (CNVs) in the aggregated data (Methods). We found that CNVs could only be readily detected and considered to be enriched in a subset of LumAcinar cells from patients with PCa, as well as in a small neuroendocrine (NE) population (Figure 6A; Figure 5—figure supplement 5B; Figure 6—figure supplements 1, 3, 4, 5, 6). Importantly, we could confidently assign LumAcinar identity to the CNV-containing tumor cells despite the transcriptomic shifts observed, due to the retained similarity of global transcriptional properties as well as specific genes among these tumor cells and adjacent benign cells (Figure 6—figure supplement 6D).

**Figure 6.**
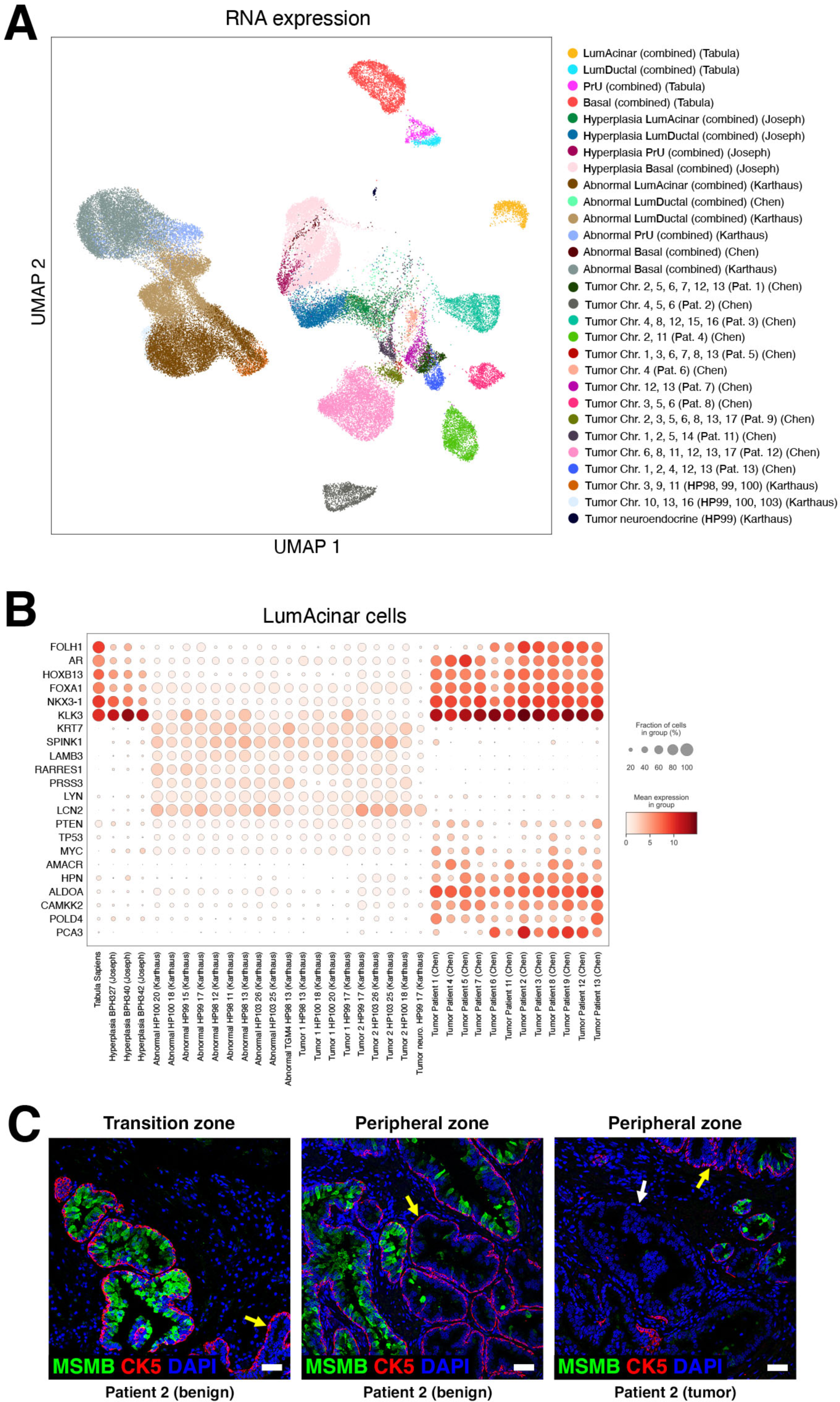
Variable CNV and gene expression patterns in LumAcinar cells from human prostate adenocarcinomas. **(A)** UMAP plot of the epithelial populations from 2 healthy prostates *(Tabula Sapiens et al., 2022)*, 3 prostates with BPH *(Joseph et al., 2020)*, and 17 prostates with treatment-naive PCa from Memorial Sloan Kettering Cancer Center *(Karthaus et al., 2020)* and Shanghai Changhai Hospital *(Chen et al., 2021)*. **(B)** Dot plot of key prostate cell type and cancer markers. **(C)** Immunofluorescence staining of varying levels of MSMB (green) in LumAcinar cells across multiple prostate zones. MSMB is expressed at high levels in normal LumAcinar cells, but is reduced or absent in abnormal acinar areas where Basal cells expressing CK5 (red) are intact (yellow arrows), and in tumor-containing areas where Basal cells are absent (white arrow).

This analysis also revealed co-occurring CNV profiles in patients from certain datasets; for example, in the Karthaus cohort ***(Karthaus et al., 2020)***, we named these tumor acinar populations “Tumor 1” (marked by CNVs on chromosomes 3, 9, and 11) and “Tumor 2” (CNVs on chromosomes 10, 13, and 16) to distinguish them from abnormal acinar populations that did not have CNV enrichment (Figure 6—figure supplement 6A-D). Interestingly, the pattern of two groups of common CNVs and expression profiles across the Karthaus cohort were not present in other datasets; for example, the Ge cohort ***(Ge et al., 2022)*** had some overlap of specific inferred CNVs, but not in the overall profiles (Figure 6—figure supplement 6). In comparison, the Chen cohort ***(Chen et al., 2021)*** displayed a more random distribution of CNVs, with no more than 2 CNVs overlapping between any patient tumors (Figure 6A, Figure 6 – supplement 4), perhaps consistent with later-stage tumors in these patients. Intriguingly, the patient 1 tumor in the Ge cohort contained multiple different clones, one with AR-high and the other with AR-low features, indicating that these different clones can coexist (Figure 6—figure supplements 5, 6).

Finally, one tumor in the Karthaus cohort contained a small neuroendocrine tumor population with a CNV profile that partially overlapped with one of the acinar tumor populations from the same region of the same tumor (Figure 6A, B; Figure 6—figure supplements 2E, 3). This observation suggests a potential common origin for these two transformed cell types.

### Transcriptomic changes in LumAcinar cells in proximity to prostate tumors

In addition to different CNV profiles, we observed fundamentally distinct features in the gene expression profiles of tumors in the Chen, Ge, and Karthaus cohorts. The tumors in the Chen cohort retained more classical acinar features, displaying increased expression for many normal LumAcinar, hyperplastic LumAcinar, and AR-responsive genes relative to tumors in the Karthaus cohort. As a result, the expression profiles of tumors in the Chen cohort overlap with those of hyperplastic acinar populations to a greater extent than those in the Karthaus cohort, as shown by signature comparisons, heatmap, and dot plot analyses (Figure 6A,B; Figure 5—figure supplement 5 A,B; Figure 6—figure supplements 3, 4). In contrast with the Chen cohort, the Karthaus tumors show significant loss of acinar features across the Tumor 1, Tumor 2, and abnormal acinar populations (as well as non-acinar populations); instead, these transformed and abnormal LumAcinar cells shifted towards PrU and LumDuctal profiles. Intriguingly, the tumor populations in the Karthaus cohort also express lower levels of selected prostate cancer-relevant markers relative to the Chen tumors, including *POLD4, AMACR,* and *CAMKK2* (Figure 6B). In comparison, the Ge cohort displayed a mixture of transcriptomic features, even within the same patient (Figure 5—figure supplement 5A,B; Figure 6—figure supplements 5, 6). These findings underscore the extent of transcriptomic variability that is already present within the treatment-naïve prostate tumors.

Notably, these gene expression changes are not limited to definitive tumor regions or transformed cell types, as we observed that a large number of acinar cells outside of the definitive tumor displayed altered transcriptomes (Figure 6—figure supplement 6A, C-G). Therefore, we performed immunofluorescence staining to validate these changes in gene expression adjacent to tumor lesions, using MSMB as a representative LumAcinar marker. We observed an apparent shift in LumAcinar gene expression close to the tumor, where the basal cell layer was disrupted, as well as at a distance from the tumor where the basal cell layer is completely intact (Figure 6C, Figure 6—figure supplement 6G; Figure 6—source data 1). These findings support the identification of “fields” of transcriptionally altered LumAcinar cells that lack CNVs and are not themselves transformed. Interestingly, this transcriptomic shift was not restricted to the peripheral zone, and transcriptomic shifts were also detected among other epithelial cell types (Figure 6—figure supplement 6D-G).

## Discussion

Single-cell analysis has revealed profound cellular and anatomical heterogeneity that has significant implications for the origin and phenotypes of prostate diseases. Our study has generated aggregated reference atlases for the human and mouse prostate, and revealed the remarkable consistency of cell types between species in tissue homeostasis as well as the plasticity displayed by prostate epithelial cells during regression and cancer.

Several notable features of the normal mouse prostate have emerged from our analyses. First, the remarkable consistency across datasets has allowed us to define 9 mouse prostate epithelial cell types and suggest an additional intermediate cell state. Second, lobe identity along the dorsal-ventral axis corresponds to the identity of a cognate distal luminal population, whereas each lobe is further divided along its proximal-distal axis into three distinct parts, corresponding to the periurethral PrU region, the proximal LumP region, and a lobe-specific distal region. Third, only luminal cell types are reliably distinct in each region along the proximal-distal axis, and thus luminal cells specifically reflect spatial identity. Finally, PrU cells have a hybrid luminal-basal identity, share gene expression with both the prostate and the urethra, and have one PrU subset displaying greater luminal features and the other more basal, which resembles the organization of the urothelium. Consequently, PrU cells have properties of a transition population at the junction of the urethra and prostate.

Our analyses of prostate regression and regeneration highlight the plasticity of both epithelial and stromal cell types, as nearly all significantly change gene expression profiles in an androgen-dependent manner. In particular, all luminal and basal epithelial populations shift after androgen deprivation towards a transcriptomic profile that resembles a PrU-like state. We speculate that this PrU-like state may mirror that of epithelial cells within the prostatic urogenital sinus and prostate epithelial buds during early organogenesis, when androgen levels are relatively low. At these early stages, both luminal and basal progenitors retain bipotent progenitor properties, and may display hybrid luminal-basal features ***(Ousset et al., 2012; Shibata et al., 2020)***. Notably, PrU cells express the highest levels of *Sca-1, Ly6d,* and other markers that have been associated with progenitor-like properties ***(Crowley et al., 2020)***. Furthermore, we have observed that treatment-naïve primary tumors often contain cells with an AR-low state resembling PrU, suggesting that PrU-like states may also recur in castration-resistant prostate cancer, when lineage plasticity leads to an increase of tumor cells with hybrid luminal-basal states ***(Chan et al., 2022)***.

Our studies have also addressed the similarity of rodent and human prostate cell populations, which has represented a long-standing question. Early histological studies suggested that the ventral lobe most closely resembled the human prostate ***(Price, 1963)***, whereas later analyses claimed that the rat dorsal lobe most closely resembled the human prostate ***(Aumuller et al., 1990; Berquin et al., 2005)***. Our meta-analysis identifies discrete LumAcinar, LumDuctal, Basal, and PrU populations in the human prostate that have transcriptomic similarities to mouse LumDist, LumP, Basal, and PrU, respectively. While the mouse LumL cells of the lateral lobe have the greatest transcriptomic similarity to human LumAcinar cells, this relationship is driven by a small number of genes, and ultimately mouse LumDist cells of all lobes generally resemble human secretory LumAcinar cells. However, the similarity of luminal cells from different zones of the human prostate to mouse luminal populations remains to be elucidated. Intriguingly, this analysis also revealed that the gene expression profiles of basal cells are very different between species, which is consistent with their unique histological and ultrastructural features such as differing basal:luminal ratios ***(El-Alfy et al., 2000)***.

Our reference atlas for the human prostate has also addressed the nomenclature for human epithelial cell types. In particular, we found that the LumDuctal and PrU populations resemble the “Club” and “Hillock” populations that were previously named due to their transcriptomic similarity to cell populations described in the lung ***(Henry et al., 2018)***. Although this is an intriguing observation, the prostate LumDuctal/Club and PrU/Hillock populations do not uniformly and specifically express the defining markers for the corresponding lung populations (*SCGB1A1* and *KRT13*, respectively). Moreover, since these prostate cell types may not have similar functions or localizations as those in the lung, we favor the use of a simpler, descriptive nomenclature and find Club- and Hillock-like cells to be subsets within the LumDuctal and PrU populations, respectively.

Our analysis of human prostate cancer has also led to several interesting findings. Notably, we only observed significant CNV alterations in LumAcinar cells and rare NE cells across independent cohorts of treatment-naïve prostate cancer ***(Chen et al., 2021; Ge et al., 2022; Hirz et al., 2023; Karthaus et al., 2020; Song et al., 2022)***, as previously noted for one of these studies ***(Hirz et al., 2023)***. This finding implies that a major cell type of origin for prostate adenocarcinoma is either a normal LumAcinar cell, or a progenitor that generates LumAcinar cells. Furthermore, primary NE prostate cancer may arise *de novo* from NE cells themselves, or from a progenitor that can give rise to NE cells, consistent with our identification of a CNV profile shared between acinar tumor cells and NE tumor cells in a patient sample. In this regard, our observation of two distinct LumAcinar cell states with neuroendocrine features in primary treatment-naïve prostate tumors may correspond to early steps in the transdifferentiation of luminal adenocarcinoma cells to neuroendocrine fates ***(Beltran et al., 2016; Chen et al., 2023; Zou et al., 2017)***. Intriguingly, both cell states as well as the NE tumor clone could be detected in the absence of androgen-deprivation therapies.

In addition, we have found that tumor adjacent benign tissue contains cells with transcriptomic alterations that are broadly present across cell types and different regional samples. Notably, although CNVs were only observed in LumAcinar and NE cells, these transcriptomic alterations were found across epithelial cell types and were validated in LumAcinar cells by immunofluorescence staining. This widespread transcriptomic reprogramming is highly suggestive of field cancerization or “field effect” in which benign tissue contains genetic or transcriptomic alterations resembling adjacent tumor tissue. Such field cancerization has been documented in many other tumor types ***(Curtius et al., 2018)***, and has been suggested in prostate cancer ***(Nonn et al., 2009)***, but is not well understood. Interestingly, this transcriptomic reprogramming was observed in multiple patient samples with hormonally-intact tumors, which suggests that widespread transcriptomic plasticity is not dependent on loss of androgen signaling.

Our findings demonstrating single-cell heterogeneity of AR signaling in treatment-naïve prostate adenocarcinomas provides deeper insights into previous studies that have classified primary tumors into subclasses with high and low AR activity ***(Mahal et al., 2018; Spratt et al., 2019)***. In particular, analyses of nearly 20,000 patient tumors analyzed by the Decipher clinical assay revealed heterogeneity of *AR* gene expression and a signature of canonical AR target genes, splitting tumors into high-AR and low-AR subsets, with the low-AR tumors displaying worse treatment response and increased expression of neuroendocrine markers ***(Spratt et al., 2019)***. These results are consistent with an independent retrospective study of over 600,000 patients, showing poorer outcomes and higher expression of neuroendocrine markers by high-grade tumors expressing low levels of prostate-specific antigen (PSA), an AR-regulated gene ***(Mahal et al., 2018)***.

Our current study indicates that this heterogeneity in AR signaling exists at the single-cell level within patient tumors, and that the previous classifications of AR-high and AR-low tumors should be further refined to reflect the heterogeneous composition of patient tumors and the possibility of tumor evolution altering the balance of AR-high and AR-low states. Given that the AR-low population transcriptionally resembles a PrU-like state, these AR-low tumor cells may display greater castration-resistance. Notably, in the mouse prostate, a transition to PrU-like expression profiles is observed in the context of regression following castration (Figure 3), with distal LumA cells displaying the most pronounced shift (Figure 3—figure supplement 3). Furthermore, PrU cells have the greatest progenitor potential among the epithelial populations in functional assays ***(Crowley et al., 2020)***, a feature that may also contribute to castration-resistance.

Finally, we have identified at least one cell state with neuroendocrine features that is associated with the AR-low population. Thus, the existence of AR-low populations within primary treatment-naïve prostate tumors raises the possibility that castration-resistance and neuroendocrine differentiation are pre-existing properties that may be selected in part by androgen-deprivation therapies. Consequently, the detection of such AR-low tumor cells in treatment-naïve tumors may represent an important step in designing precision therapies for primary prostate cancer.

## Methods

### Key Resources Table

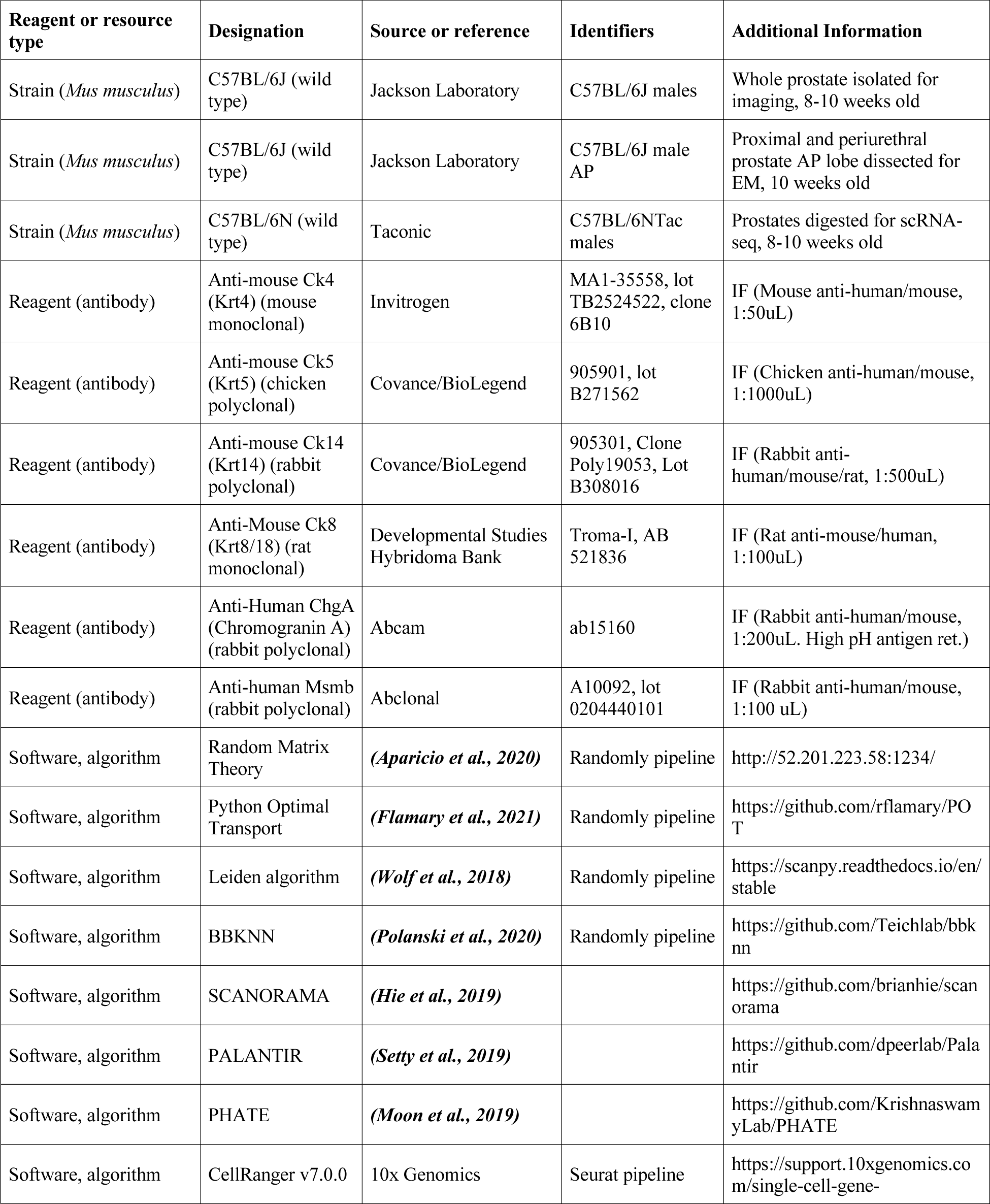

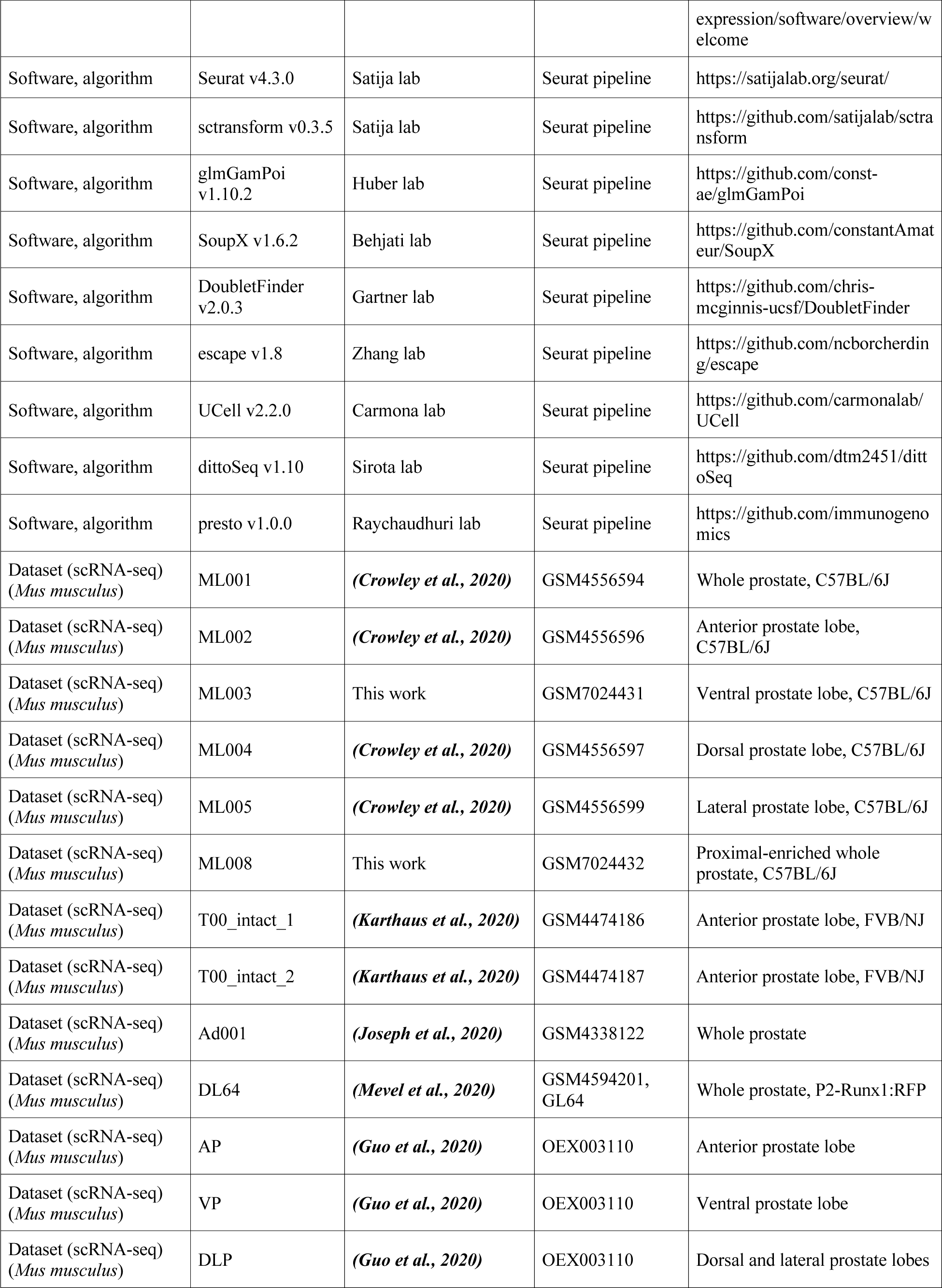

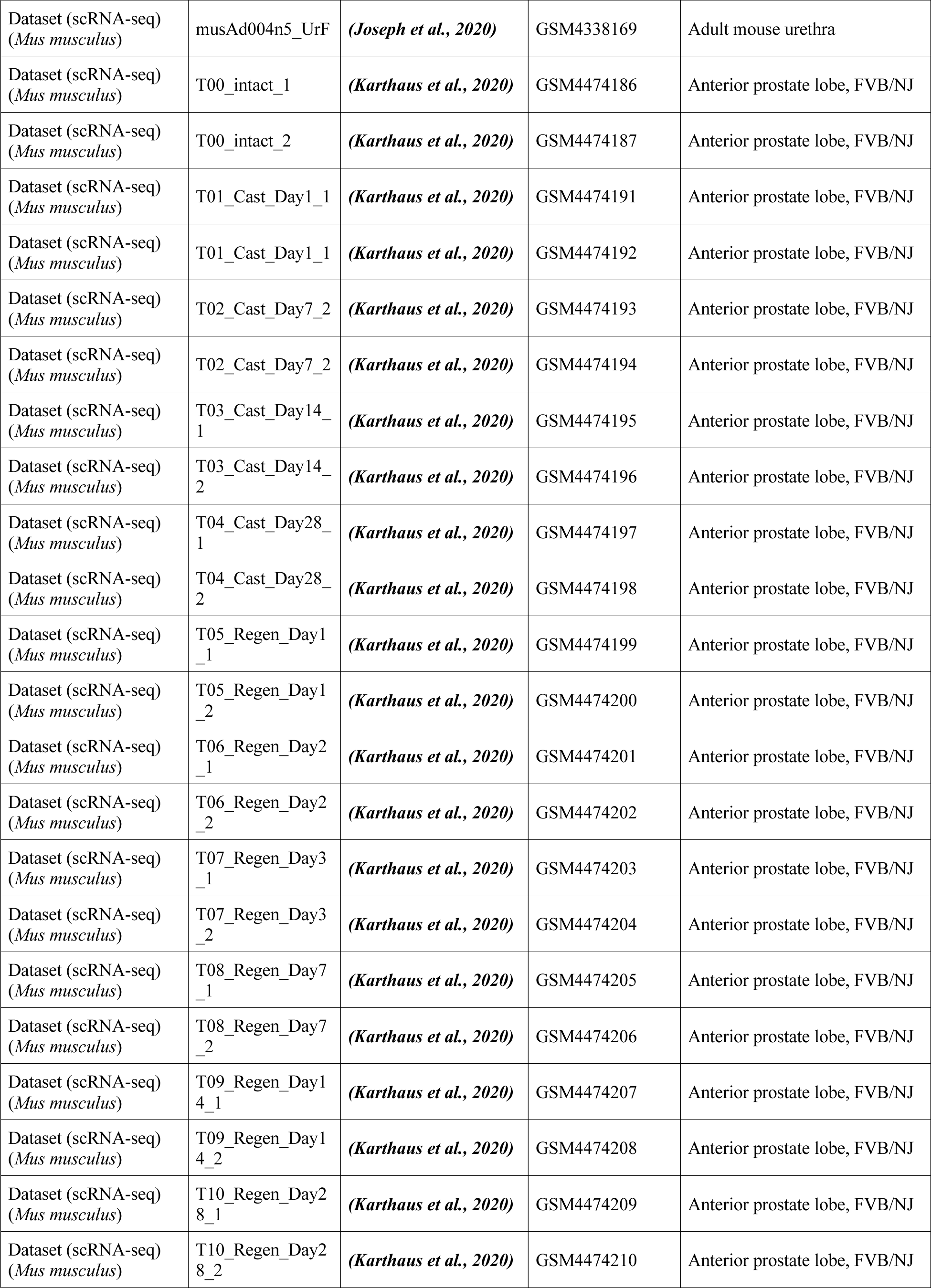

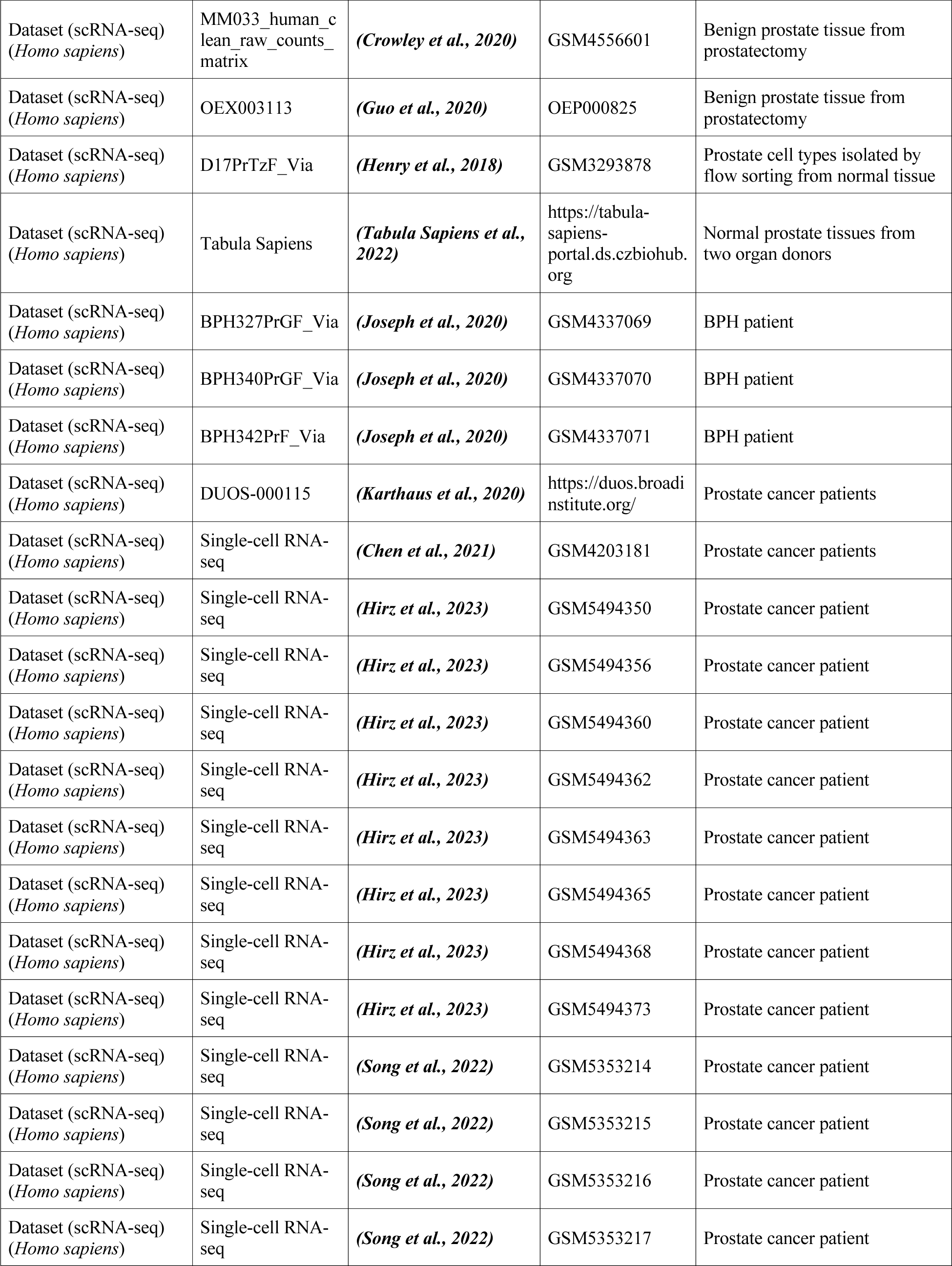

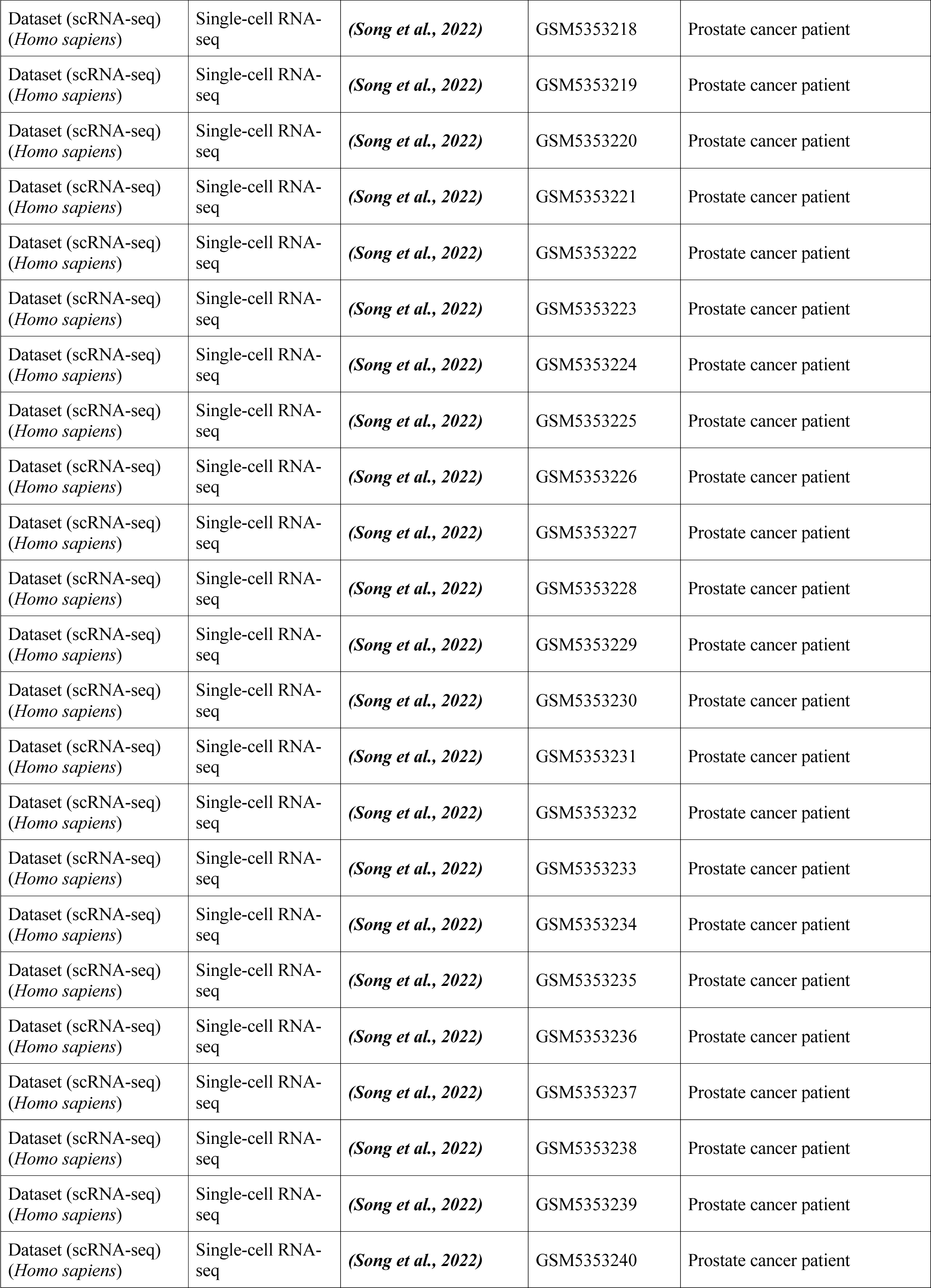

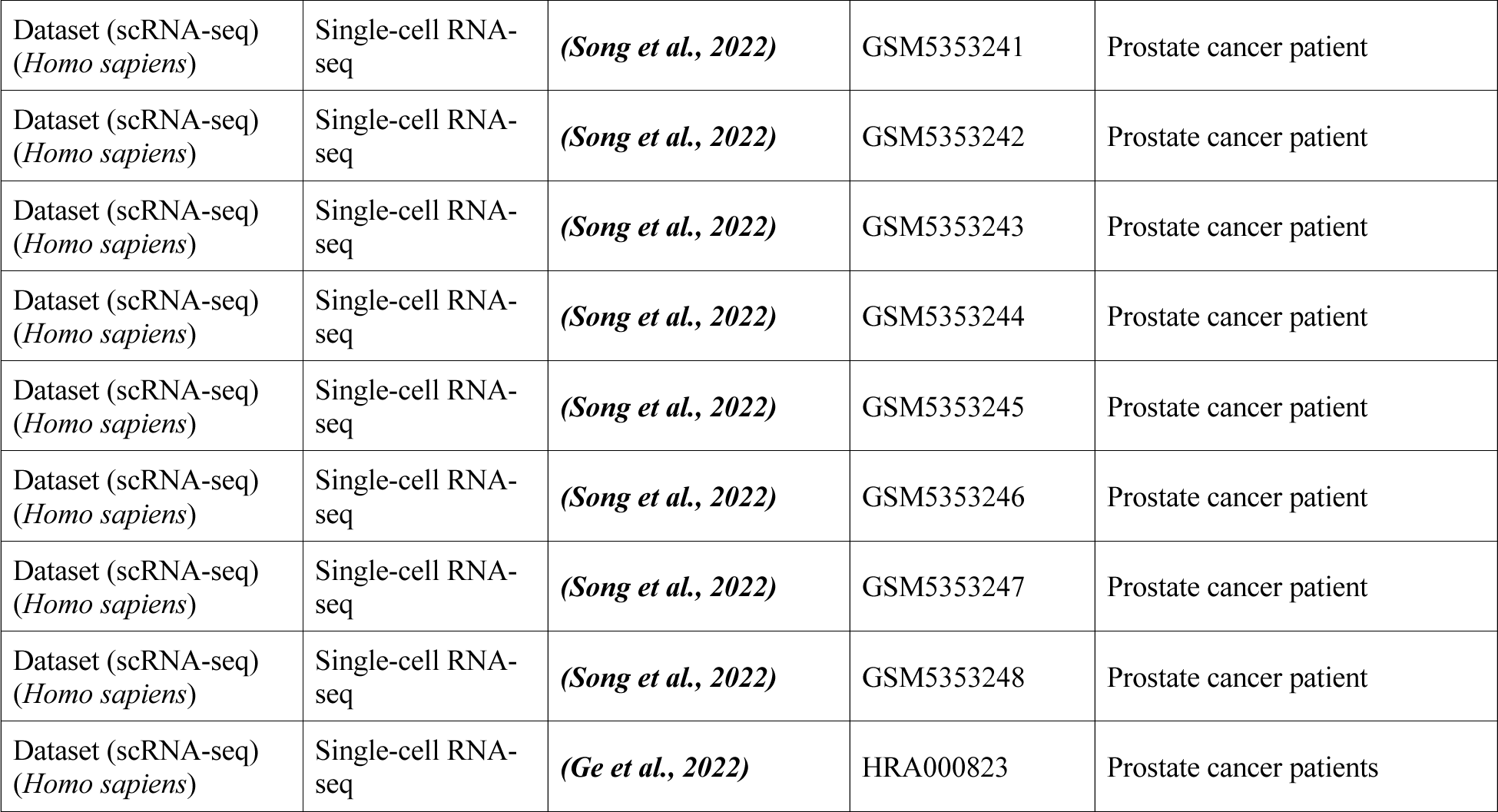

### Continuing analyses

Mouse strains and genotyping, isolation of mouse prostate tissue, dissociation of mouse prostate tissue, and prostate single-cell RNA-sequencing was carried out as previously described ***(Crowley et al., 2020)***. Additionally, immunofluorescent imaging was performed as previously described ***(Crowley et al., 2020)***.

### Isolation of mouse prostate tissue

Tissue was isolated from wild type C57BL6/N (C57BL/6NTac, 8-10 week old) mice to generate the new datasets described here. For the ventral prostate (VP) lobe dataset (ML003/ GSM7024431), the entire extent of the VP lobes was dissected from one male mouse at 8 weeks of age, from the distal tips to the proximal end within the rhabdosphincter. For the proximal and periurethral prostate dataset (ML008/GSM7024432), a proximally-enriched region was dissected from 3 male mice, 10 weeks of age. The rhabdosphincters were removed, and prostate tissue was collected from the periurethral junction with the urethra on one end (including minimal surrounding urethra), to 1-2 mm beyond the proximal:distal boundary on the other end (to include some distal cells). Additionally, a tiny region of proximal seminal vesicle (SV) was dissected from 2 mice to include in the sample after removal of secretions. These samples were processed as previously described ***(Crowley et al., 2020)***. All animal studies were approved by and conducted according to standards set by the Columbia University Irving Medical Center (CUIMC) Institutional Animal Care and Use Committee (IACUC).

### Electron microscopy

Prostate tissue was taken from a C57BL/6 mouse at 8 weeks of age. An approximately 2 mm region of the AP lobe within the rhabdosphincter near the periurethral-proximal boundary was micro-dissected. The sample was fixed, processed, sectioned, and imaged as previously described ***(Crowley et al., 2020)***.

### Mouse prostate analyses

Two separate pipelines were used in parallel to analyze available mouse prostate datasets: Seurat, and randomly. The Seurat pipeline is summarized below. The Randomly pipeline is similar to what was previously published ***(Crowley et al., 2020)***, with detailed description below.

### Seurat scRNA-seq analysis pipeline

FASTQ files for all datasets were either already in house or downloaded from the Short Read Archive (SRA) or the National Omics Data Encyclopedia (NODE). FASTQ files were then aligned and quantified using CellRanger v7.0.0. All scRNA-seq counts were corrected for ambient RNA using the SoupX package. The cleaned counts were then converted into Seurat objects using the Seurat package. Cells with high mitochondrial DNA content, low gene detection, and/or high RNA counts were filtered out to enrich for live single cells. Datasets were normalized, and their variances were stabilized with SCTransform. Cell doublets were then computationally detected and filtered out using DoubletFinder. All individual datasets were then merged into a new Seurat object, their original counts normalized, and their variance stabilized using SCTransform.

To produce an integrated dataset, integration anchors were calculated, and the datasets were then integrated using the reciprocal principal component analysis (RPCA) reduction in Seurat. A new PCA reduction and UMAP reduction were then generated using the integrated dataset. For first-pass cluster calling, neighbors and clusters were determined using Seurat at a resolution of 0.8 and a Louvain algorithm with multilevel refinement. These clusters were then minimally manually adjusted to reflect physical anatomy and marker expression previously validated by immunofluorescence staining. Gene set enrichment analysis was performed using the escape package with the “UCell” method and Hallmark mouse gene sets provided by MSigDB. Visualizations of the gene set enrichment analysis were performed using the dittoSeq package.

Mouse prostate population signatures were generated in Seurat dataset with the wilcoxauc function of the presto package on the aggregated dataset. Signature genes with the most globally distinguished expression patterns were determined by applying a filter to collect only genes with an AUC ≥ 0.75 and an adjusted p-value ≤ 0.05. This was performed for each distinct cell type/cluster, as well as for informative subgroups (such as all distal luminal cells versus proximal luminal cells).

### Randomly scRNA-seq analysis pipeline

Randomly analyses were conducted as previously described ***(Crowley et al., 2020)***. Sequencing data were aligned and quantified using the Cell Ranger Single-Cell Software Suite (v.2.1.1) with either the GRCm38 mouse or the GRCh38 human reference genomes. There are 4 major steps: 1) filtering the raw sequencing data expression matrix, 2) correcting for batch effects using Seurat and processing with Randomly (http://52.201.223.58:1234/) ***(Aparicio et al., 2020)***, 3) clustering data using the Leiden algorithm (https://scanpy.readthedocs.io/en/stable), and 4) dimensional reduction for visualizations such as t-SNE and UMAP plots (included in Randomly package). Departures from our previous methods will be summarized below.

### Filtering the expression matrix

Cell-gene matrices were pre-processed by filtering cells with less than 500 genes detected. We also removed cells whose proportion of transcripts derived from mitochondrially encoded genes was greater than 10%. The expression matrices were normalized by *log*_2_(1 + *TPM*), where ‘() is transcripts per million.

### Random Matrix Theory application to denoise scRNAseq

Random Matrix Theory (RMT) was first introduced by Wishart in 1928, but the mathematical foundations of RMT were developed by the theoretical physicist Dyson in the 1960s when he was describing heavy atomic nuclei energy levels. A key feature of RMT is universality, namely the insensitivity of certain statistical properties to variations of the probability distribution used to generate the random matrix. This property provides a unified and universal way to analyze single-cell data ***(Aparicio et al., 2020)*** and we previously used this method to describe new cell populations in prostate ***(Crowley et al., 2020)***.

The RMT strategy relies on the fact that single-cell datasets show a threefold structure: a random matrix, a sparsity-induced signal, and a biological signal. Indeed, 95% or more of the single-cell expression matrix is compatible with being a random matrix ***(Aparicio et al., 2020)***. This could be understood as if the dataset is showing cells whose expression is randomly sampled from a given distribution in approximately 95% of the matrix inputs. In single-cell datasets, sparsity is also a key feature, as it can generate a fake signal that after removal increases the quality and performance of clustering in prostate scRNA-seq analyses, and led to identification of the novel PrU population ***(Crowley et al., 2020)***. From an operative point of view, the presence of localized eigenvectors related with sparsity implies the existence of an undesired (fake) signal.

### Clustering

Clustering was performed using the Leiden algorithm, as implemented in ***(Polanski et al., 2020; Wolf et al., 2018)***. The determination of the optimal number of clusters relied on the mean silhouette score. Specifically, we conducted a series of clustering analyses across various Leiden resolutions (the clustering parameter) and calculated the mean silhouette score for each scenario. We established a relationship between the mean silhouette score, acting as a function of the Leiden resolution, and the respective number of clusters for each case (see Figure 1—figure supplement 1 in ***(Crowley et al., 2020)***). We selected the absolute maximum of this curve, and took the corresponding number of clusters. In certain instances, sub-clustering specific clusters proved beneficial. The process involved repeating the described procedure for a designated cluster. Sub-clustering was particularly valuable for unraveling immune populations or differentiating between vas deferens and seminal vesicle populations. The robustness of sub-clustering was verified through supervised plotting of known genes associated with the aforementioned populations.

### Batch effect correction and data integration

In Figures 1A and 4A, the samples corresponding to mice (Figure 1) and humans (Figure 3) were aggregated, employing the BBKNN method ***(Polanski et al., 2020)*** with default parameters. In Figure 5A, the data for luminal acinar cells from normal prostate, BPH and PCa were integrated using SCANORAMA ***(Hie et al., 2019)***, using default parameters.

### Differential expression analysis

The genes highlighted and presented in the dot-plots were chosen using a strategy based on differential expression. These selected genes underwent a t-test (one group vs. all others) with a corrected p-value (Benjamini-Hochberg correction) below 0.01. Additionally, as a secondary threshold for selection, these genes were required to display expression in a minimum of 60% of cells within the target population and in less than 25% of cells for all other populations.

Gene signatures for each human epithelial population were generated using a similar approach. All of the most differentially expressed genes for each population were selected that had corrected p-values of ≤ 0.05, with the secondary requirement of expression in a minimum of 60% of the cells within the target population and a maximum of 25% of the cells for other populations.

### Representations and visualizations

To visualize the single-cell clusters, we performed dimensional reduction to two dimensions through t-distributed Stochastic Neighbor Embedding (t-SNE) and Uniform Manifold Approximation and Projection (UMAP) representations. Default parameters were utilized for both techniques: a learning rate of 1000, perplexity of 30, and early exaggeration of 12 for t-SNE; for UMAP, we set the number of neighbors to 15 and minimum distance to 0.3. Visualizations using t-SNE, such as dot-plots or ridge-plots, were carried out using the visualization functions from the Randomly public package in ***(Aparicio et al., 2020)***, and the visualization functions within SCANPY in ***(Polanski et al., 2020; Wolf et al., 2018)***.

In Figure 5A, a two-dimensional visualization was performed using PHATE ***(Moon et al., 2019)***, depicting the luminal acinar cells from 2 normal prostates ***(Tabula Sapiens et al., 2022)***, 3 prostates with BPH ***(Joseph et al., 2020)***, and 46 prostates with PCa from Memorial Sloan Kettering Cancer Center ***(Karthaus et al., 2020)***, University of California, San Francisco ***(Song et al., 2022)***, Massachusetts General Hospital ***(Hirz et al., 2023)***, Peking University Third Hospital ***(Ge et al., 2022)***, and Shanghai Changhai Hospital ***(Chen et al., 2021)***, using default parameters. The data were integrated using SCANORAMA ***(Hie et al., 2019)***.

### Pseudotime analysis

To depict the developmental trajectory and cellular fate of the aggregated luminal acinar cells depicted in Figure 5A, we employed PALANTIR ***(Setty et al., 2019)*** for the detection of single-cell trajectories in pseudotime.

### Cell type score

For the analyses of Figure 3A and B, we constructed a “cell type score” to quantify the changes in transcriptomic profile for each cell type, based on the genes that are most specific and differentially expressed among the Basal, LumA, LumP, Mes1, Mes2, myofibroblast, smooth muscle and vascular endothelial populations. The cell type score was generated by assessing the mean expression of a specific set of differentially expressed genes that effectively characterize each population. The chosen genes for the cell type score underwent a t-test (one group vs. all others) with a corrected p-value (Benjamini-Hochberg correction) less than 0.01. Additionally, these genes were required to be expressed in at least 60% of cells within the target population and in fewer than 25% of cells for all other populations. To construct the cell type score, the mean expression of differentially expressed genes for each population was compared at each timepoint during regression. This was followed by division by the mean expression of the genes in the normal tissue before castration, and the resulting values were normalized to a scale of 1-100.

### Identification of tumor cells

We applied InferCNV ***(Tickle et al., 2019)*** to the scRNA-seq datasets to discern malignant epithelial cells exhibiting genomic instability. Epithelial cells classified as non-malignant based on copy number alterations (CNV) via inferCNV may represent authentic benign cells or transformed cells lacking identifiable CNVs through scRNA-seq inference. The analytical approach involved initial examination of each patient sample independently, employing denoising and clustering transcriptomic analyses as detailed above, to identify cell populations akin to those in the human consensus atlas. Subsequently, inferCNV was executed for each patient sample within the same cohort to pinpoint cell populations with CNVs.

To serve as control populations in inferCNV, various non-epithelial cell types were employed, along with external controls sourced from Tabula Sapiens ***(Tabula Sapiens et al., 2022)***. For the Karthaus cohort ***(Karthaus et al., 2020)***, where only epithelial cells were identified, Tabula Sapiens external controls were exclusively utilized as a reference population. In contrast, for the remaining cohorts we used mesenchymal populations detected in these samples and populations from Tabula Sapiens as controls.

For each tumor, the CNV matrix obtained from inferCNV was presented in a heatmap, displaying previously identified populations with copy number alterations, using the standard/default scale, and applying the option of hierarchical clustering to visualize the heatmap. Further insights into copy number differences were derived through dimensional reduction of the CNV matrix analysis, employing PCA with its first 20 principal components. This dissimilarity was illustrated in a UMAP, revealing that clusters lacking clear CNVs, which include internal controls such as mesenchymal cells, tended to aggregate together, whereas clusters with distinct CNVs formed separate groupings (Figure 6—figure supplements 3, 4, 5, and 6).

Epithelial populations devoid of CNVs but observed in cancer patients were labeled as “abnormal”, based on changes in their transcriptomic profiles (Figure 6—figure supplement 6). This categorization applied to LumAcinar cells lacking CNVs as well as non-acinar epithelial populations in Figure 6A and Figure 6—figure supplement 6-6. We note that the approach utilized for identifying tumor cells via inferCNV has intrinsic limitations, given its basis on inference from scRNA-seq data, and that many alterations such as mutations that might be present in tumor cells would not be captured by the techniques analyzed in this study.

### Assessing population similarity using optimal transport theory

We employed Optimal Transport ***(Kolouri et al., 2017; Villani, 2003)*** to evaluate the transcriptomic similarity between cell types, as done previously ***(Crowley et al., 2020)***. The Wasserstein-1 distance serves as a metric for phenotypic distance among cell populations, defined as a distance function between probability distributions in a measurable metric space. Conceptually, the Wasserstein-1 distance aligns with the earth mover’s distance, wherein probability distributions are envisioned as piles of dirt, and the cost of transforming one pile into another corresponds to the Wasserstein distance. We employed this approach to compare normal tissue with each time point during regression and regeneration (Figure 3—figure supplement 3), using the aggregated mouse prostate dataset from Figure 1A as a reference.

A similar methodology was applied to assess the similarity between mouse and human prostate epithelial populations (Figure 4—figure supplement 4D). In this case, optimal transport and Wasserstein distance were utilized to compare the aggregated mouse dataset with Tabula Sapiens. Initially, we identified orthologous gene pairs, separately normalized mouse and human datasets using *log*_2_(1 + *TPM*), filtered out genes with an average expression less than 0.1 for human or mouse, and merged the corresponding mouse and human datasets. Employing RMT to eliminate sparsity-induced signals, we selected genes with biological signals in the shared mouse/human dataset. Subsequently, we calculated the Wasserstein distance in the common space between mouse and human, visualizing these distances through a set of nested heatmaps (Figure 4—figure supplement 4D). Finally, in Figure 5D, Wasserstein distance was utilized to compare the phenotype of cancer cell states depicted in Figure 5A with normal tissue cell types.

## Data availability

Single-cell RNA-sequencing data from this study have been deposited in the Gene Expression Omnibus (GEO) under the accession number GSE224452. The publicly available datasets used for analyses are listed in the Key Resources Table.

## Acknowledgements

We would like to thank Cory Abate-Shen, Brett Carver, and Charles Sawyers for helpful discussions on this work, and thank Erin Bush and Peter Sims for assistance with single-cell sequencing. This work utilized the Columbia Genomics and High Throughput Screening Shared Resource as well as the Molecular Pathology Shared Resource of the Herbert Irving Comprehensive Cancer Center, which is supported in part by the Cancer Center Support Grant P30CA013696. For assistance with electron microscopy, we thank Alice Liang, Chris Petzold, and Joseph Sall at the New York University Langone Health DART Microscopy Lab, which is partially funded by NYU Cancer Center Support Grant P30CA016087 and by S10OD019974. We also want to thank Teresa Rosa for her assistance in organizing the project. These studies were supported by NIH grants R01CA238005 (M.M.S.), R01CA251527 (M.M.S.), P01CA265768 (M.M.S), U01CA261822 (R.R. and M.M.S.), R35 CA253126 (R.R.), U01 CA243073 (R.R.), by SU2C Convergence 3.14 (R.R.), by the Prostate Cancer Foundation (M.M.S.), and by fellowships from the NIH (F32CA261152; J.C.) and the National Science Foundation DGE 16-44869 (L.C.).

## Author contributions

Conceptualization: L.A., L.C., R.R., and M.M.S.; Methodology: L.A., L.C.; Formal analysis: L.A., L.C., J.R.C.; Investigation and experiments: L.C.; Resources: C.J.L., H.H.; Data curation: L.A., L.C., and J.R.C.; Writing (original draft): L.C.; Writing (review and editing): L.C., M.M.S., L.A., H.H., and R.R.; Supervision: R.R. and M.M.S.; Funding acquisition: L.C., J.R.C., R.R., and M.M.S.

---

**Figure 1—figure supplement 1.**
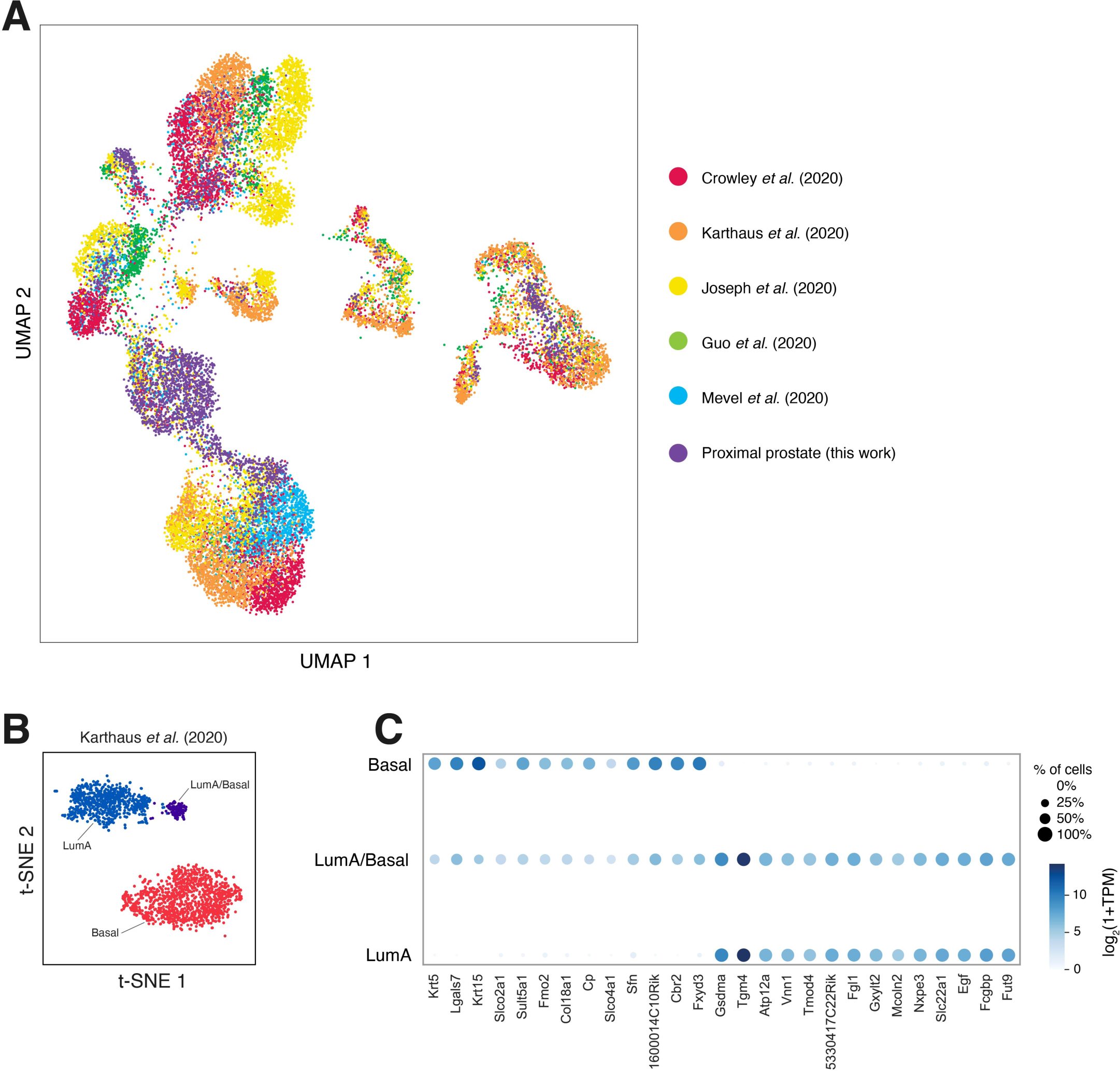
Supporting data for Figure 1A. **(A)** Alternative visualization of the aggregated dataset in Figure 1A to show source dataset for each population. **(B,C)** Additional subclustering suggests an intermediate LumA/Basal cell state in a specific dataset ***(Karthaus et al., 2020)***, as shown by a UMAP plot **(B)** and dot plot of marker expression **(C)**.

**Figure 1—figure supplement 2.**
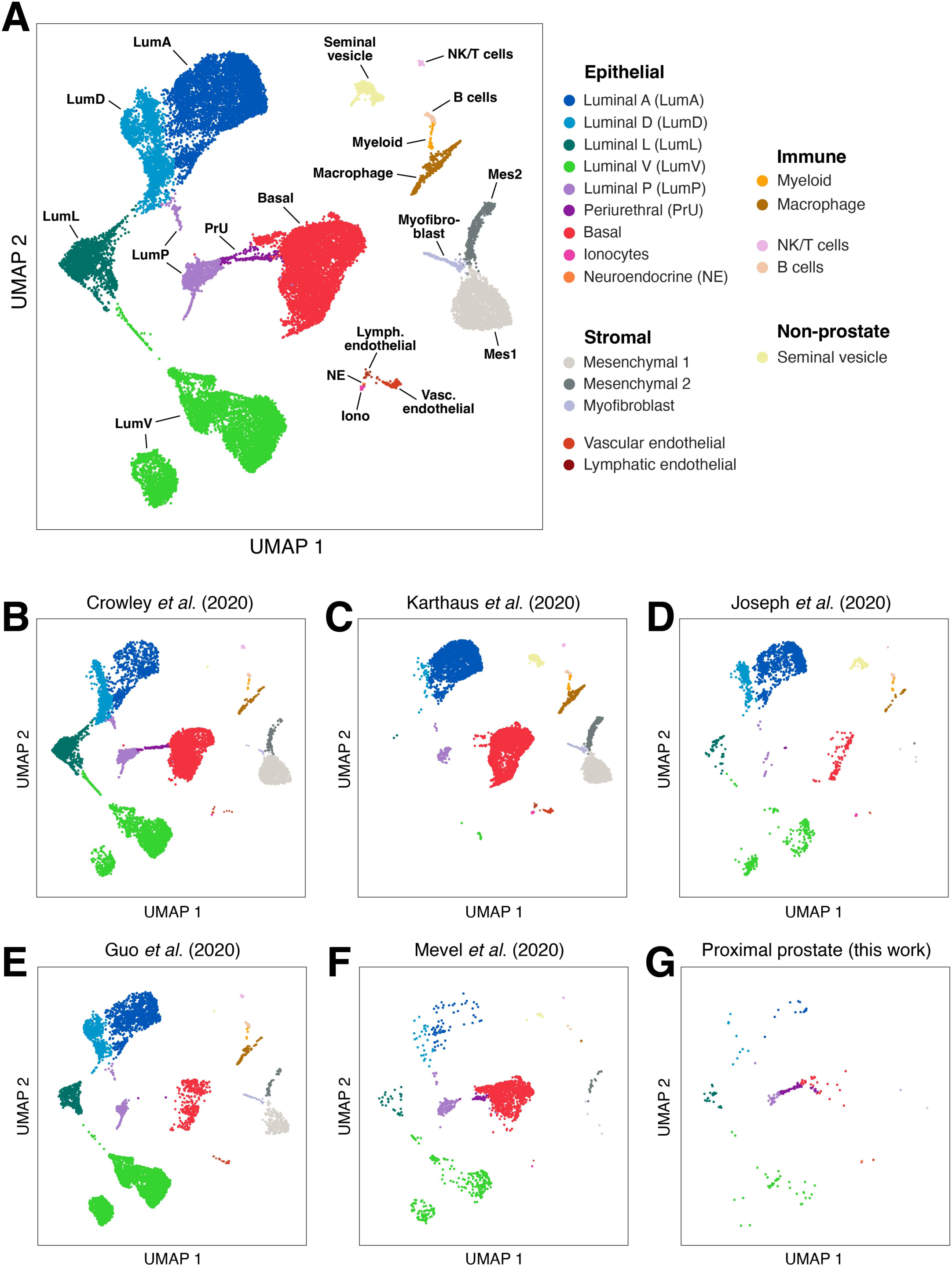
Additional subclustering of specific cell populations. **(A,B)** The immune compartment from ***(Karthaus et al., 2020)*** contains at least 5 distinct myeloid populations **(A)**; dot plot shows distinguishing markers **(B)**. **(C,D)** The immune compartment from ***(Joseph et al., 2020)*** contains at least 4 distinct myeloid populations **(C)**; dot plot showing distinguishing markers **(D)**. **(E,F)** The stromal compartment from ***(Karthaus et al., 2020)*** contains 4 distinct populations with potential further subdivision of the Mes2 population. **(G,H)** The non-prostate cells from ***(Karthaus et al., 2020)*** reveal potential proximal/distal subdivision of the seminal vesicles. **(I,J)** The ionocytes from ***(Karthaus et al., 2020)*** show potential heterogeneity as well as presence in both prostate and seminal vesicle tissue.

**Figure 1—figure supplement 3.**
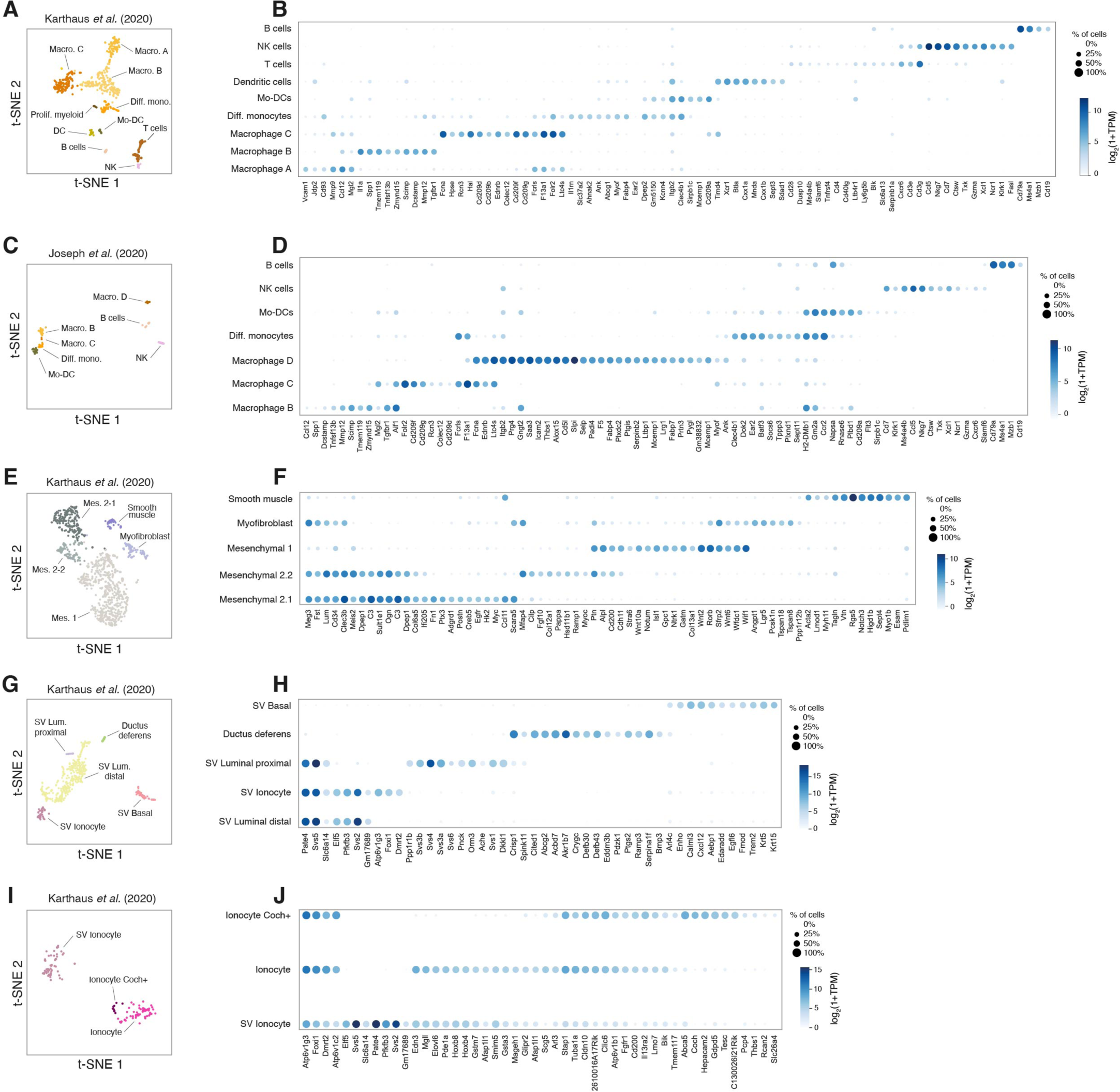
Reference plots of mouse prostate scRNA-seq data generated by the Seurat pipeline. Shown are plots arranged as in Figure 1, revealing 9 epithelial populations, 5 stromal populations, and at least 4 immune populations across multiple datasets. This approach reproduced all epithelial and stromal populations; however, it did not subcluster some immune populations, so these were labeled as combined populations. B-G) Manuscript sources of the datasets that compose the aggregate UMAP, showing intermixing of cells from different mouse strains and processing methods within population clusters.

**Figure 1—figure supplement 4.**
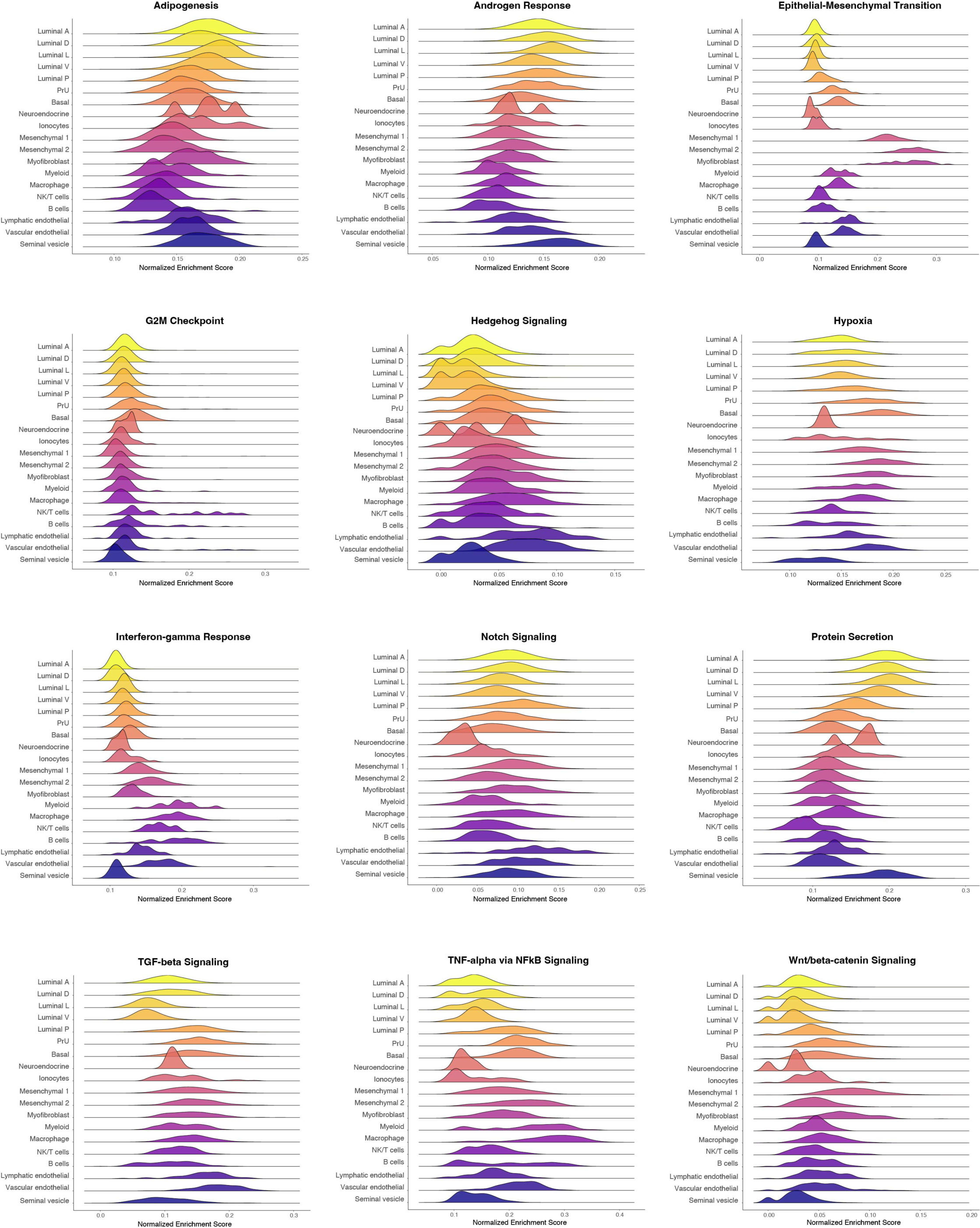
Gene Set Enrichment Analysis of the mouse prostate populations using Hallmark Pathways. Pathway analysis of curated gene expression signatures shows the normalized expression of gene sets involved in different processes by cell type. Analyses for each pathway were performed on aggregated mouse prostate data in the Seurat pipeline.

**Figure 2—figure supplement 1.**
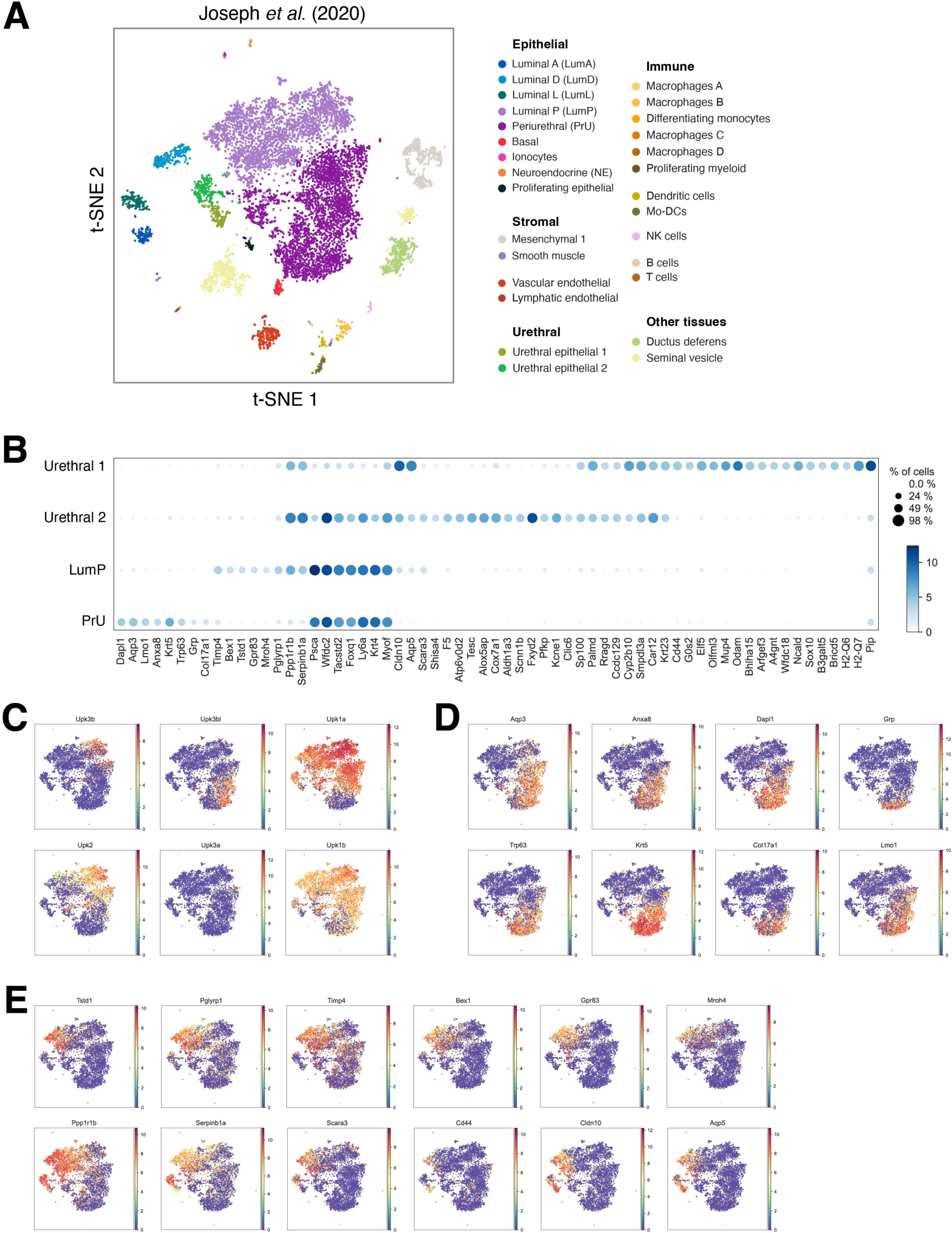
Meta-analysis of a published dataset for the mouse periurethral region. **(A)** UMAP visualization of dataset from ***(Joseph et al., 2020)*** shows distal, proximal, periurethral, basal, neuroendocrine, and ionocyte cells, mesenchymal and immune populations, and several non-prostate populations such as the urethra. **(B)** Dot plot of differentially expressed markers between LumP (mixed), PrU (mixed), and urethra-only populations. Although they are visibly distinct and physically separated, both the prostate PrU and LumP populations have gene expression overlap with similar urethral populations. **(C-E)** Genes with enriched expression in the urothelium **(C)**, prostatic PrU **(D)**, and prostatic LumP **(E)**.

**Figure 3—figure supplement 1.**
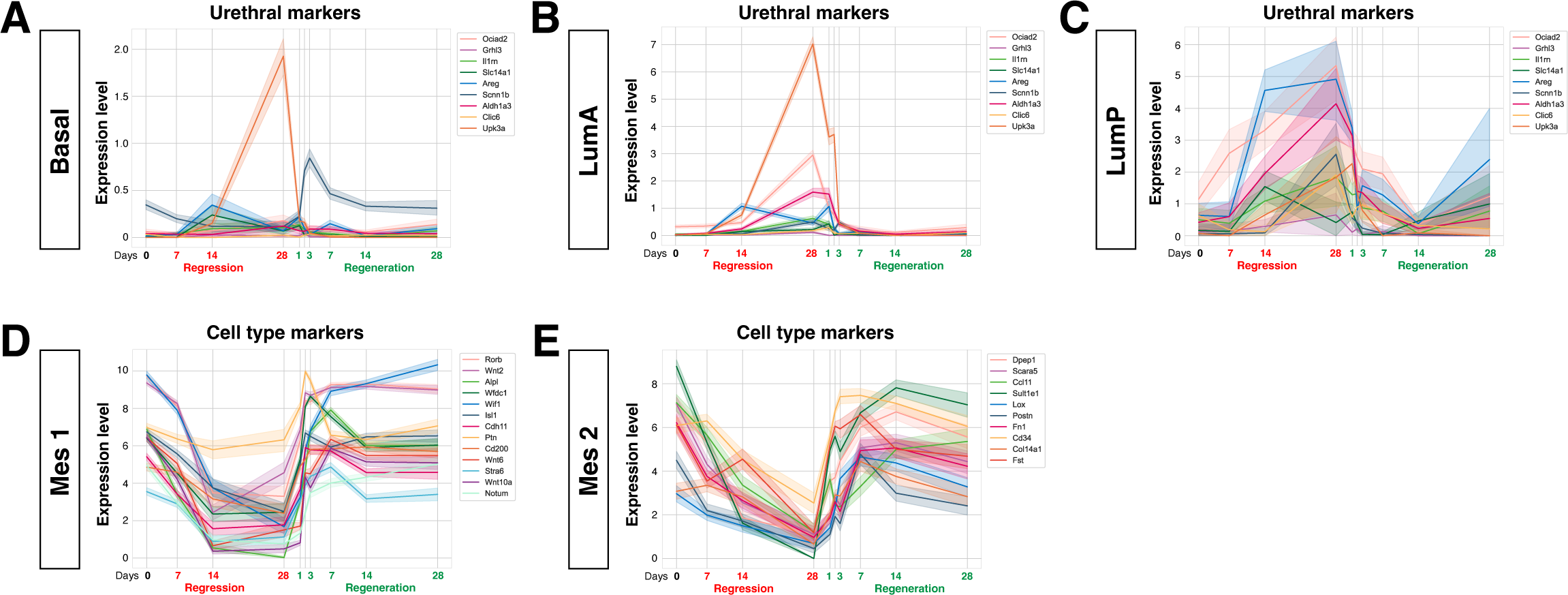
Gene expression changes during prostate regression and regeneration. Line plots generated from meta-analysis of a regression-regeneration time course based on ***(Karthaus et al., 2020)***, where the line indicates average gene expression across the population and the bar indicates the CI (+/- 95%). **(A-C)** Genes that are enriched in urethral and not PrU cells, in Basal **(A)**, LumA **(B)**, and LumP **(C)** cells. LumA and Basal cells generally did not gain expression of urothelial markers, with the exception of Upk3a, which is specific to the urethra in C57BL/6 mice, whereas LumP cells gained expression of a subset of urothelial markers in addition to the PrU markers. **(D,E)** Expression of distinguishing markers for the Mes 1 (D) and Mes 2 (E) fibroblast populations over the regression and regeneration time course.

**Figure 3—figure supplement 2.**
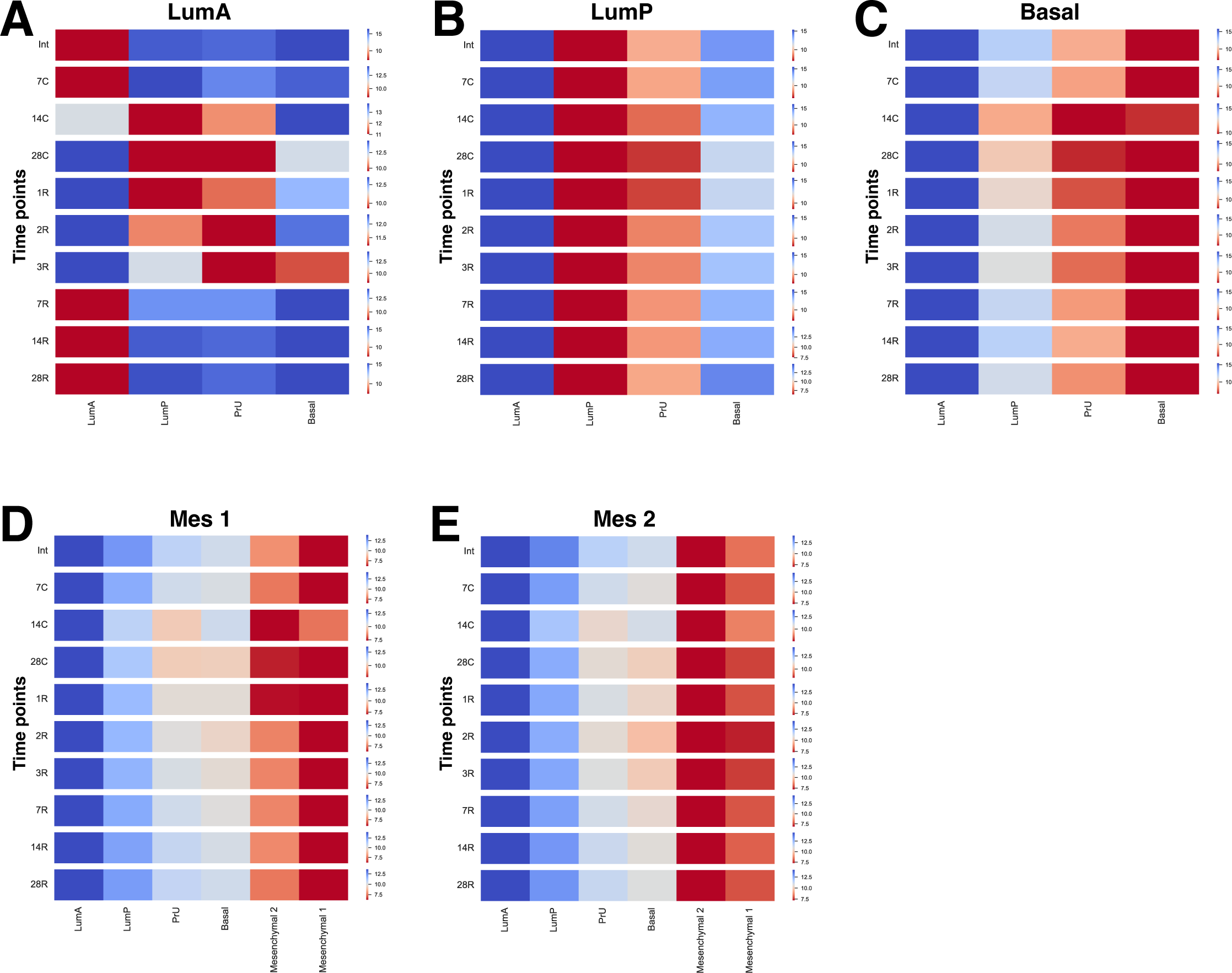
Heatmaps of total gene expression profiles for individual cell populations during prostate regression and regeneration. Profiles of LumA **(A)**, LumP **(B)**, Basal **(C),** as well as Mes 1 **(D)** and Mes 2 **(E)** based on data from *(Karthaus et al., 2020)* are compared to the normal gene expression profiles from the Tabula Sapiens. Measurements are given in Wasserstein distance, a metric of effort required to convert one profile into the other; red indicates greater similarity, and blue indicates less similarity between populations.

**Figure 4—figure supplement 1.**
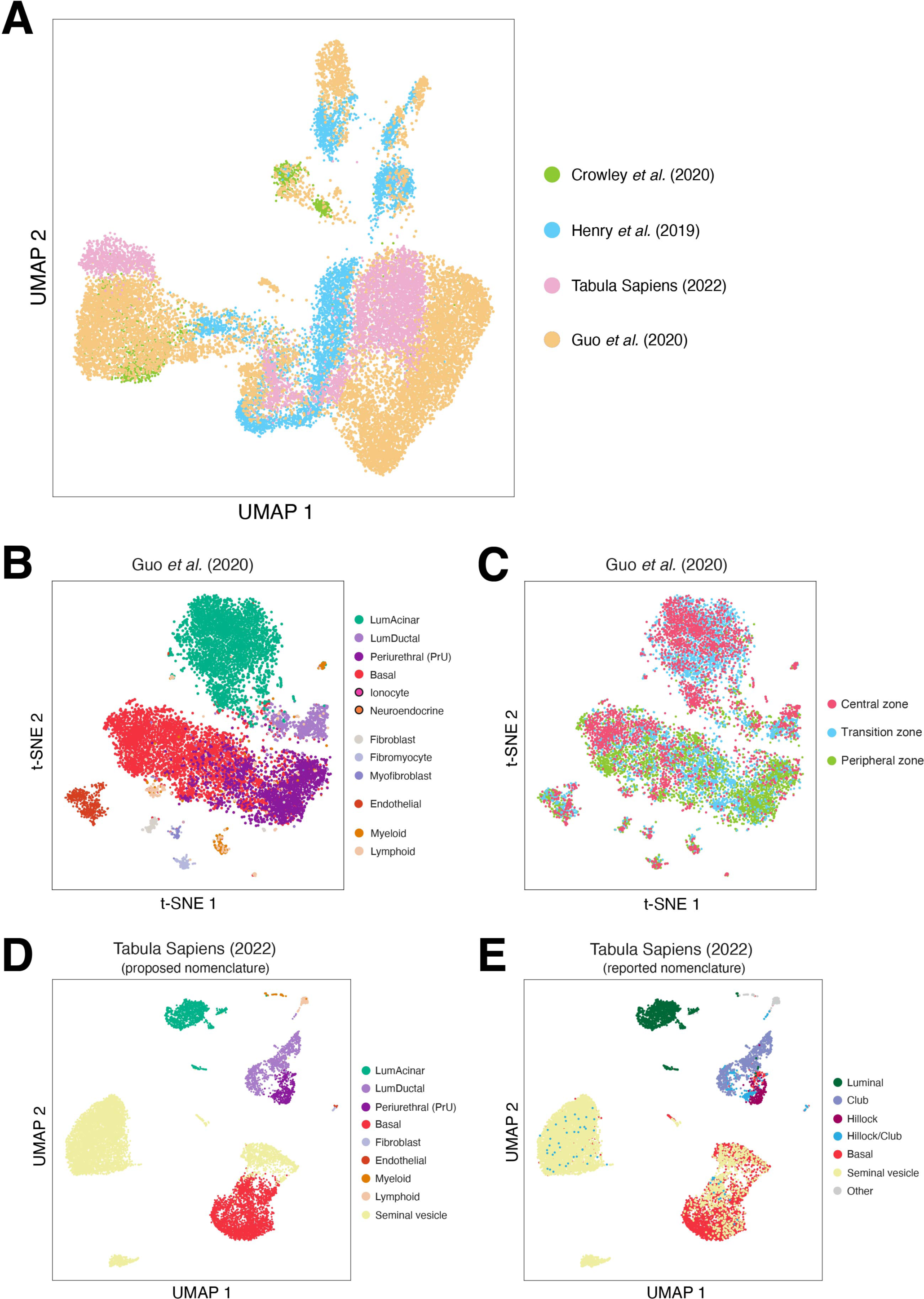
Additional analysis of the normal human prostate atlas. **(A)** Alternative visualization of the aggregated dataset in Figure 4A showing the source datasets. **(B,C)** UMAP of the data from ***(Guo et al., 2020)*** using our descriptive nomenclature as shown in Figure 4C **(B)**, and the zonal identity of each cell **(C)**. Although each large epithelial population contains cells from each zone, proportionally fewer of the LumAcinar cells were from the peripheral zone. **(D, E)** Human prostate nomenclature shown for the Tabula Sapiens normal prostate dataset ***(Tabula Sapiens et al., 2022)***, comparing the proposed nomenclature **(D)** to the nomenclature from ***(Henry et al., 2018)* (E)**.

**Figure 5—figure supplement 1.**
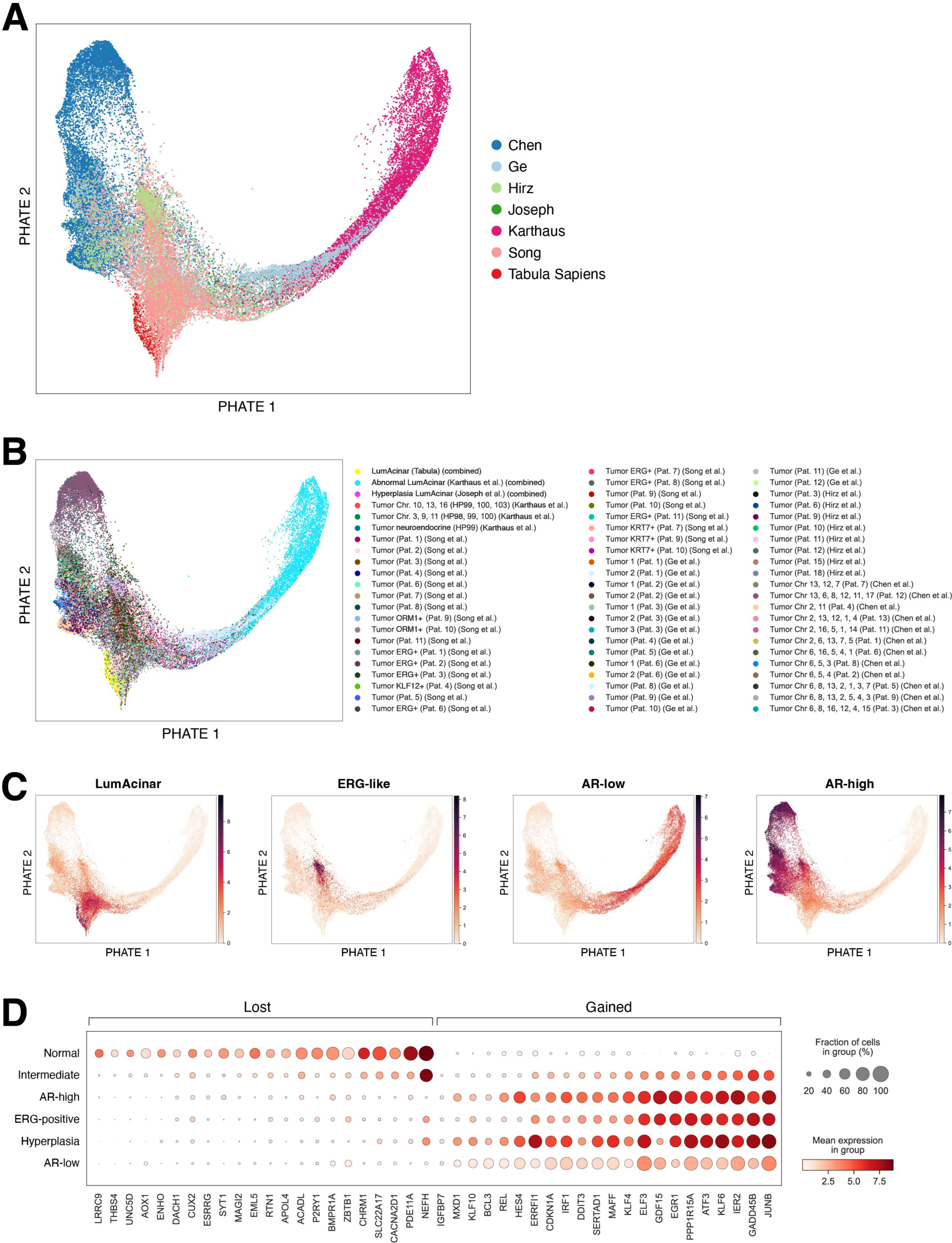
Gene expression profiles and cohort details for LumAcinar cells from normal and transformed prostates. **(A-C)** PHATE plots of LumAcinar cells corresponding to Figure 5A. **(A)** Data contribution from the 6 cohorts *(Chen et al., 2021; Ge et al., 2022; Hirz et al., 2023; Joseph et al., 2020; Karthaus et al., 2020; Song et al., 2022; Tabula Sapiens et al., 2022)*. **(B)** Disease state, subgroup, and tumor samples for the patients within each cohort. CNV analyses are detailed in the Methods and in Figure 6. (**C)** Average expression of the gene sets associated with normal and transformed LumAcinar cell states. Transformed and abnormal cells exhibit heterogeneous profiles that divide into those with high androgen receptor signaling (high-AR), ERG-positive, or low AR signaling. **(D)** Dot plot displaying selected genes with differential expression between normal and transformed LumAcinar cells, divided into those lost (left) or gained (right) during transformation.

**Figure 5—figure supplement 2.**
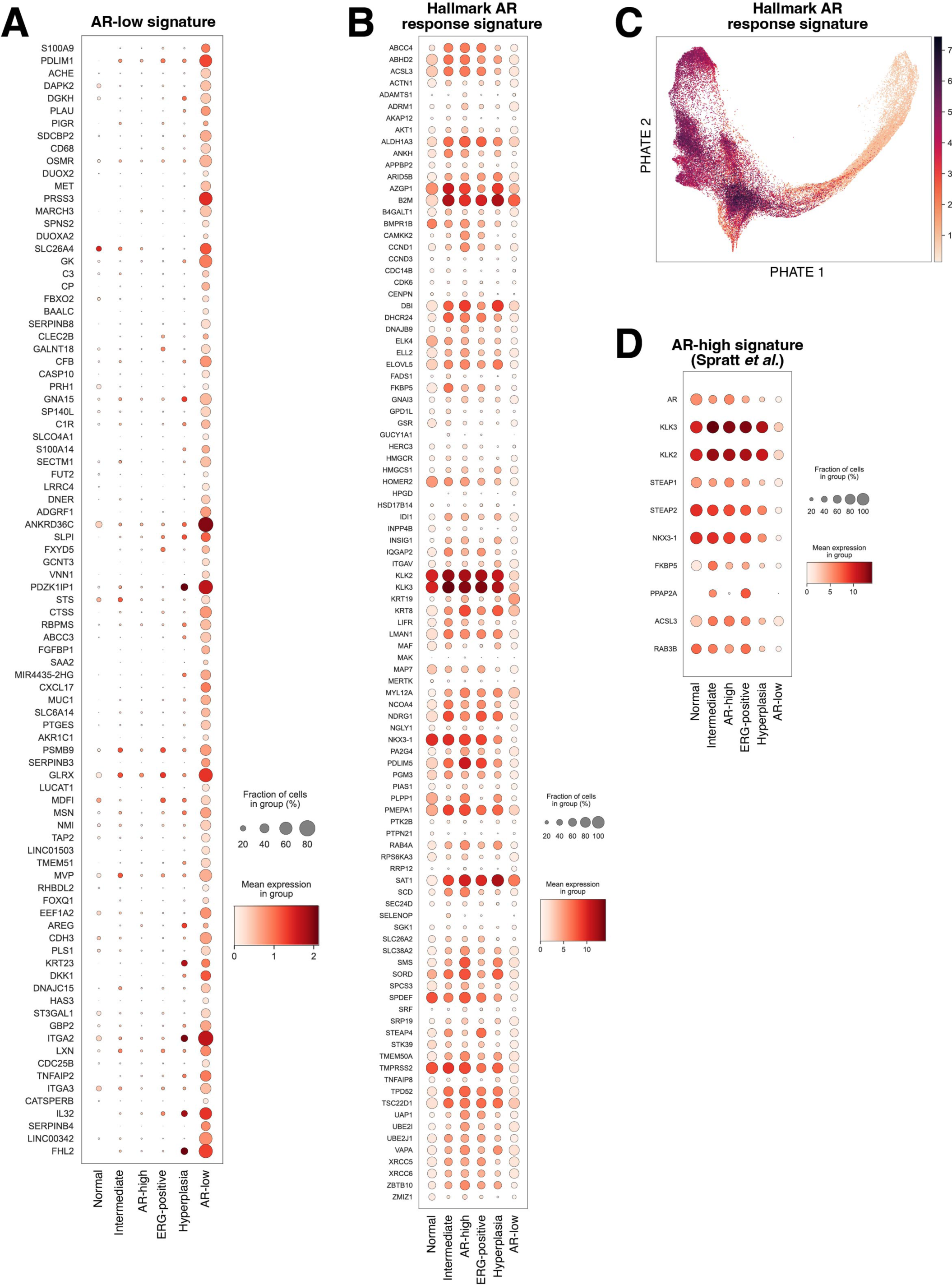
Gene expression profiles for AR-high and AR-low transformed LumAcinar cells. **(A)** Dot plot of the 90 most differently expressed genes in LumAcinar cells of the AR-low subgroup, corresponding to the AR-low signature. **(B)** Dot plot of the Hallmark Androgen Response Signature genes. **(C)** PHATE plot of the mean expression of the Hallmark Androgen Response gene set. **(D)** Dot plot of the AR-high signature from ***(Spratt et al., 2019)***. Both signatures depict a broad negative correlation with the AR-low signature genes in **(A)**.

**Figure 5—figure supplement 3.**
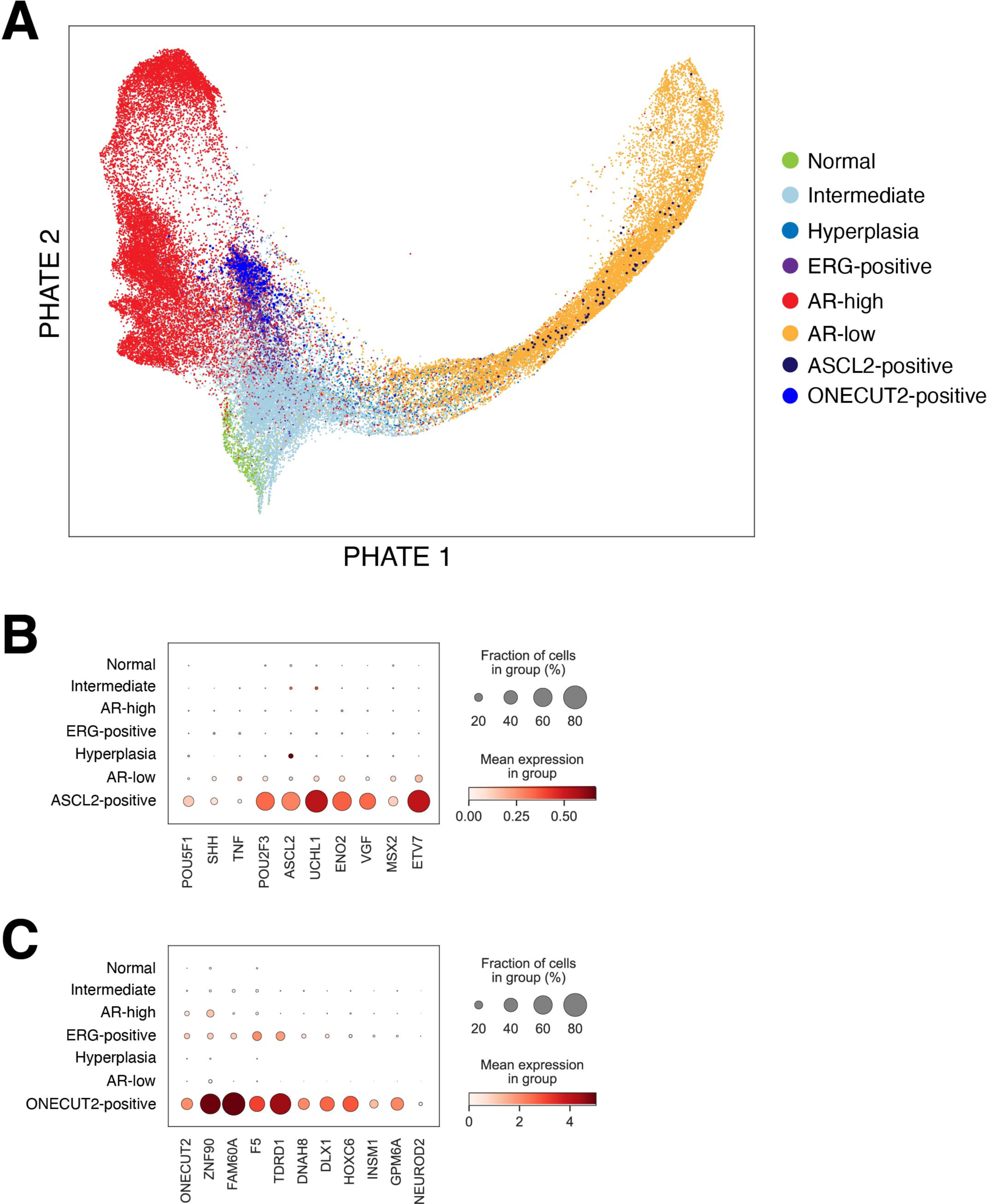
Markers of neuroendocrine differentiation in tumor and abnormal acinar cells. **(A)** PHATE plot highlighting the location of two distinct subsets of cells expressing neuroendocrine markers were identified through manual curation based on collective expression of genes associated with neuroendocrine differentiation within the entire group. The ASCL2-positive subset predominantly overlaps with the AR-low population, and the ONECUT2-positive subset aligns with the AR-high and ERG-positive populations. **(B, C)** Dot plots depicting selected genes that characterize the ASCL2-positive **(B)** and ONECUT2-positive **(C)** subsets.

**Figure 6—figure supplement 1.**
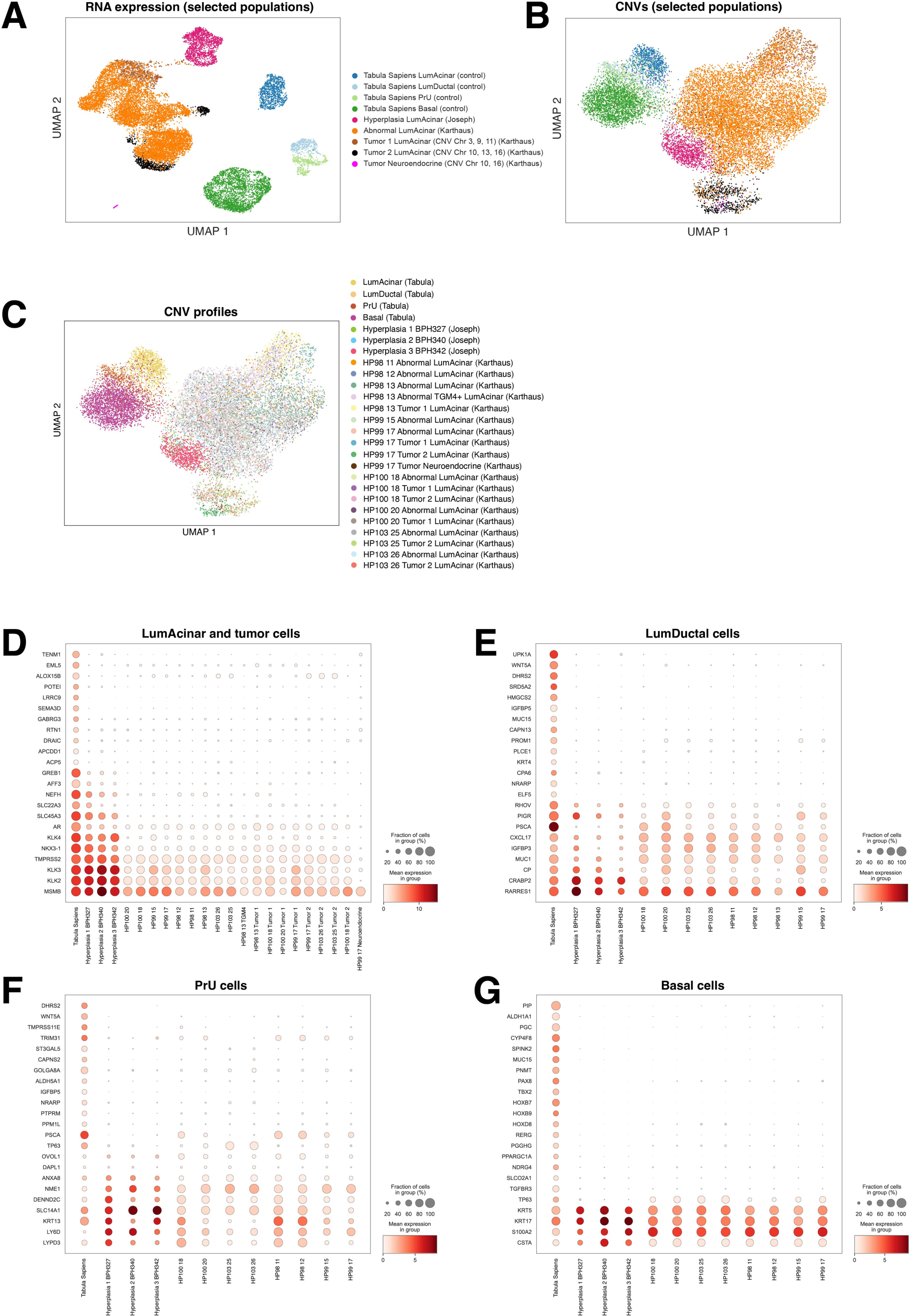
Meta-analysis of human prostate scRNA-seq data across disease states reveals altered gene expression profiles and a LumAcinar bias of CNVs. **(A,B)** UMAP plots of RNA expression **(A)** and CNV profiles **(B)** of epithelial populations in 2 normal prostates *(Tabula Sapiens et al., 2022)*, 3 patients with BPH *(Joseph et al., 2020)*, and 4 patients with treatment-naive PCa *(Karthaus et al., 2020)*. To identify tumor populations, inferCNV was used to detect CNVs across the disease samples, normalized against all epithelial populations of the normal prostate from Tabula Sapiens *(Tabula Sapiens et al., 2022)*, and clusters with CNVs were labeled as definitive tumors. **(C)** UMAP plot of CNV profiles for each acinar cluster shows that hyperplastic cells have similar CNV profiles to abnormal acinar cells while definitive tumor cells are distinct. **(D-G)** Dot plots of differentially expressed genes, for the LumAcinar and NE populations **(D)**, LumDuctal **(E)**, PrU **(F)**, and Basal cells **(G)** across these datasets. For LumAcinar and NE cells in which tumor populations were detected, populations are listed in order of CNV enrichment. Population names match the names used in the source datasets, including spatial regions for each sample from *(Karthaus et al., 2020)*; additional classification was added in **(D)** to distinguish tumor acinar cells from abnormal acinar cells.

**Figure 6—figure supplement 2.**
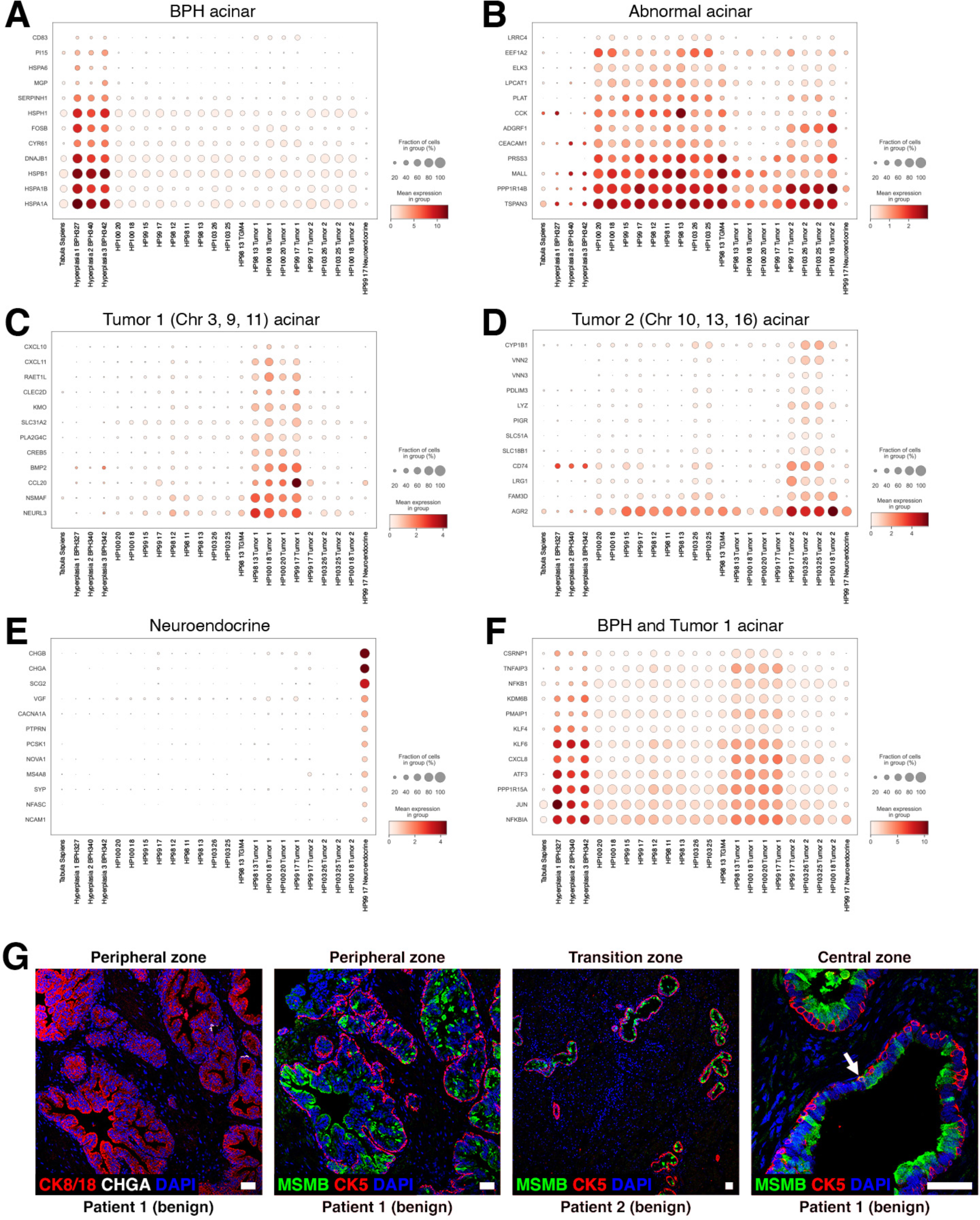
Marker expression for the LumAcinar and NE populations across disease states. **(A-F)** Dot plots of specific markers for BPH acinar cells **(A)**, abnormal acinar cells **(B)**, Tumor CNV group 1 acinar cells **(C)**, Tumor CNV group 2 acinar cells **(D)**, Neuroendocrine tumor cells **(E)**, and both BPH and Tumor 1 acinar cells **(F)**. Note that BPH acinar cells uniquely express genes for many heat shock proteins, and that abnormal acinar cells generally express the same genes as definitive tumor acinar cells, even when these cells were isolated from different regions of the prostate. **(G)** Immunofluorescence analysis of cell type and LumAcinar markers. *(Left)* MSMB is expressed in CK8/18-positive LumAcinar cells in the peripheral zone, and is absent where Basal cells are still intact, independent of proximity to NE cells. *(Middle right)* Reduced MSMB expression in LumAcinar cells can also be observed in the transition zone, which is less frequently a site of prostate cancer. *(Right)* Rare example of MSMB expression in a CK5+MSMB+ intermediate-like cell.

**Figure 6—figure supplement 3.**
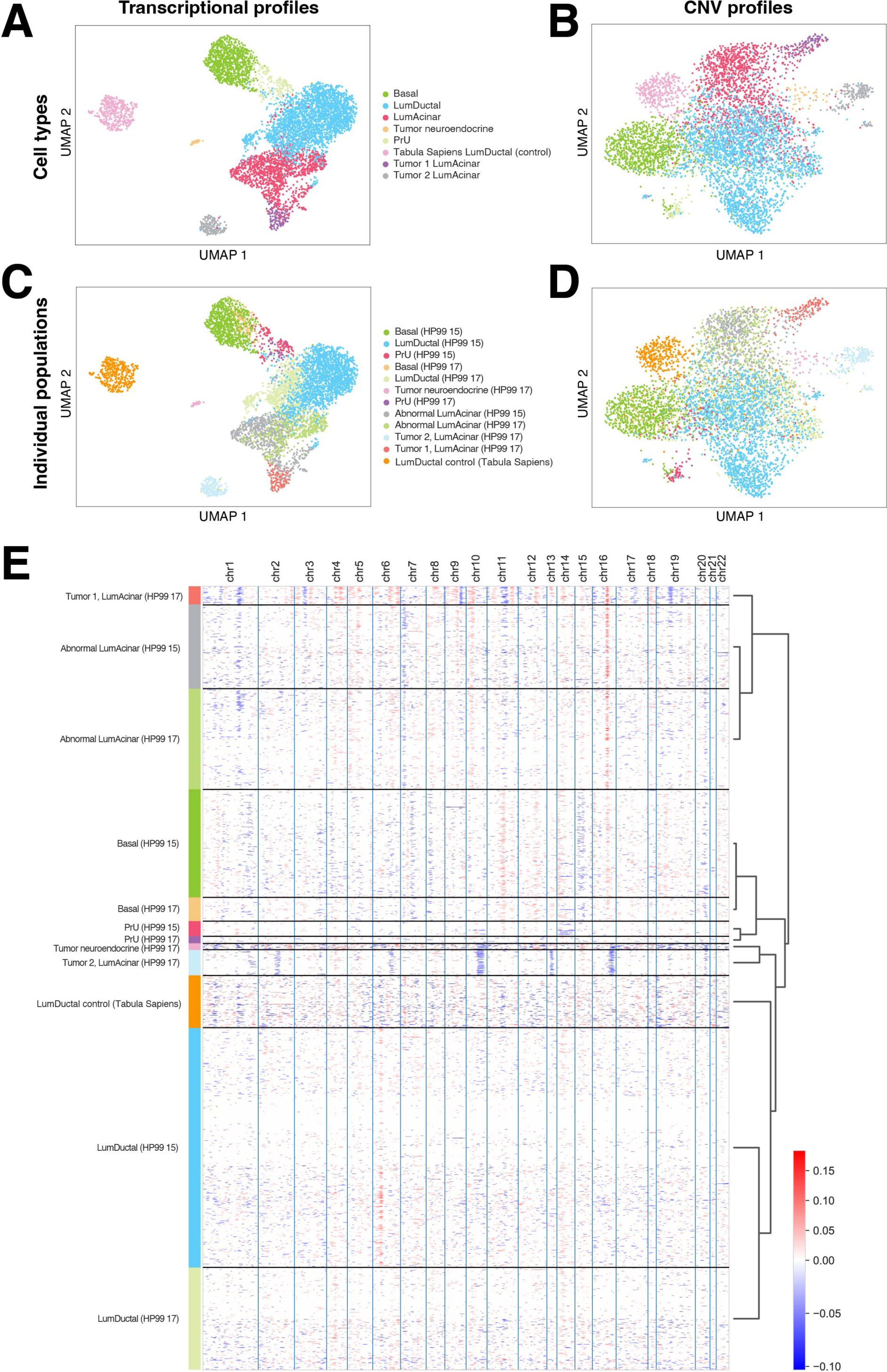
InferCNV methodology to distinguish tumor and abnormal cell populations. **(A-D)** UMAP plots of gene expression **(A,C)** and CNVs **(B,D)** in prostate cell populations from patient HP99 *(Karthaus et al., 2020)*. **(A,B)** show general cell populations and the Tabula Sapiens Luminal Ductal cells (used for CNV normalization), and **(C, D)** show specific populations detected by the RMT pipeline. **(E)** Heatmap of detected CNVs across sections 15 and 17 taken from HP99, as well as the control LumDuctal cells *(Tabula Sapiens et al., 2022)* used for normalization. A population was determined to be a definitive tumor population if over 25% of the cells within the population were enriched for the same CNV over a control. HP99 has a Tumor 1 group population (CNVs on chromosomes 3, 9 & 11) and additional patient-specific CNVs, a Tumor 2 group population (CNVs on chromosomes 10, 13 & 16) and additional CNVs, and a NE Tumor population (CNVs on chromosomes 10 & 16). This *de novo* NE tumor population shares several CNVs with an acinar Tumor 2 population from the same region. Definitive tumor populations for HP99 were all detected in section 17, though transcriptomic changes could be detected throughout populations in both sections 15 and 17, which have unique cell type compositions.

**Figure 6—figure supplement 4.**
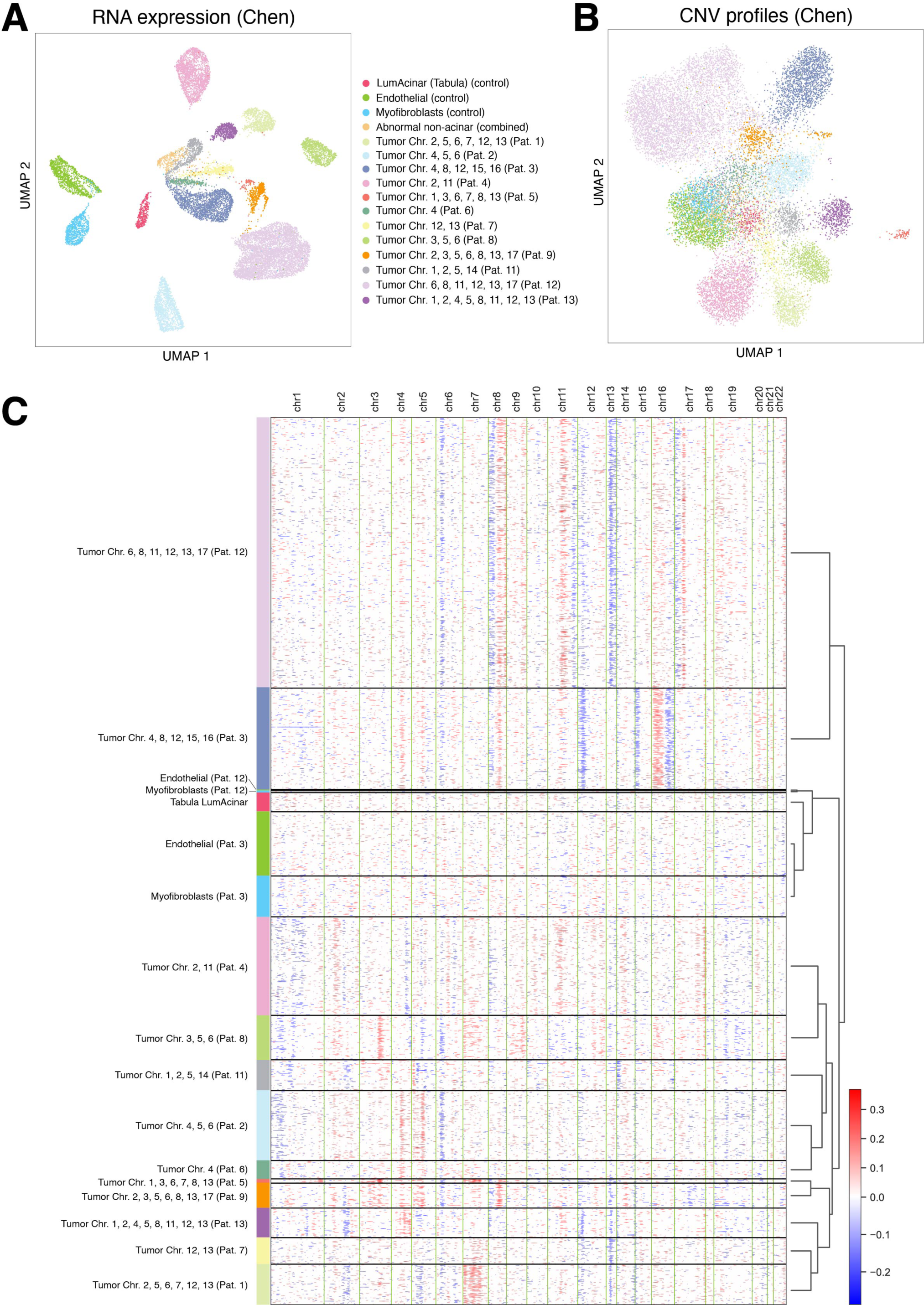
InferCNV analysis of prostate tumors from the Chen cohort. **(A,B)** UMAP plots of gene expression **(A)** and CNVs **(B)** for the cell populations from *(Chen et al., 2021)* and a normalization control from *(Tabula Sapiens et al., 2022)*. **(C)** Heatmap of CNV enrichment across pooled patient samples and control populations. Data were pooled and CNVs were normalized as a group to both internal mesenchymal populations and external normal LumAcinar cells from Tabula Sapiens. All definitive tumor populations are LumAcinar, and this cohort has completely variable CNV patterns between patients. Of note, one patient (SC173) in the Chen cohort may have undergone androgen-deprivation therapy, but it is unclear whether this sample was sequenced and/or included in our analysis.

**Figure 6—figure supplement 5.**
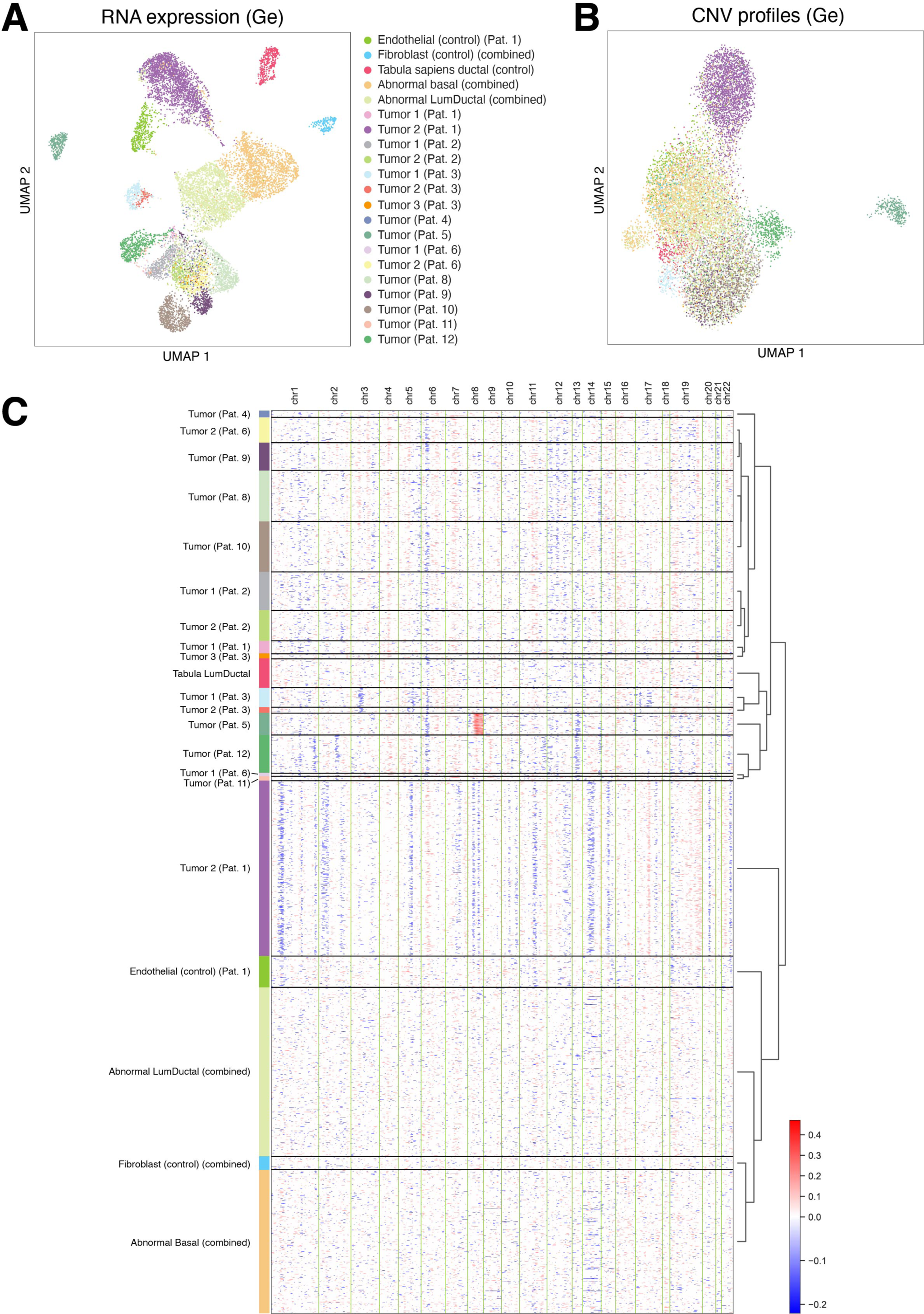
InferCNV analysis of prostate tumors from the Ge cohort. **(A,B)** UMAP plots of gene expression **(A)** and CNVs **(B)** for the cell populations from 12 PCa patients ***(Ge et al., 2022)*** and a normalization control from ***(Tabula Sapiens et al., 2022)***. **(C)** Heatmap of CNV enrichment across the patient samples. Data were pooled and CNVs were normalized as a group to multiple internal mesenchymal and epithelial populations as well as external normal LumAcinar cells from Tabula Sapiens. This cohort has CNV patterns that are observed in several patients; further, many patients had multiple tumor populations with distinct sets of CNVs, which we have labeled numerically. All definitive tumor populations are LumAcinar.

**Figure 6—figure supplement 6.**
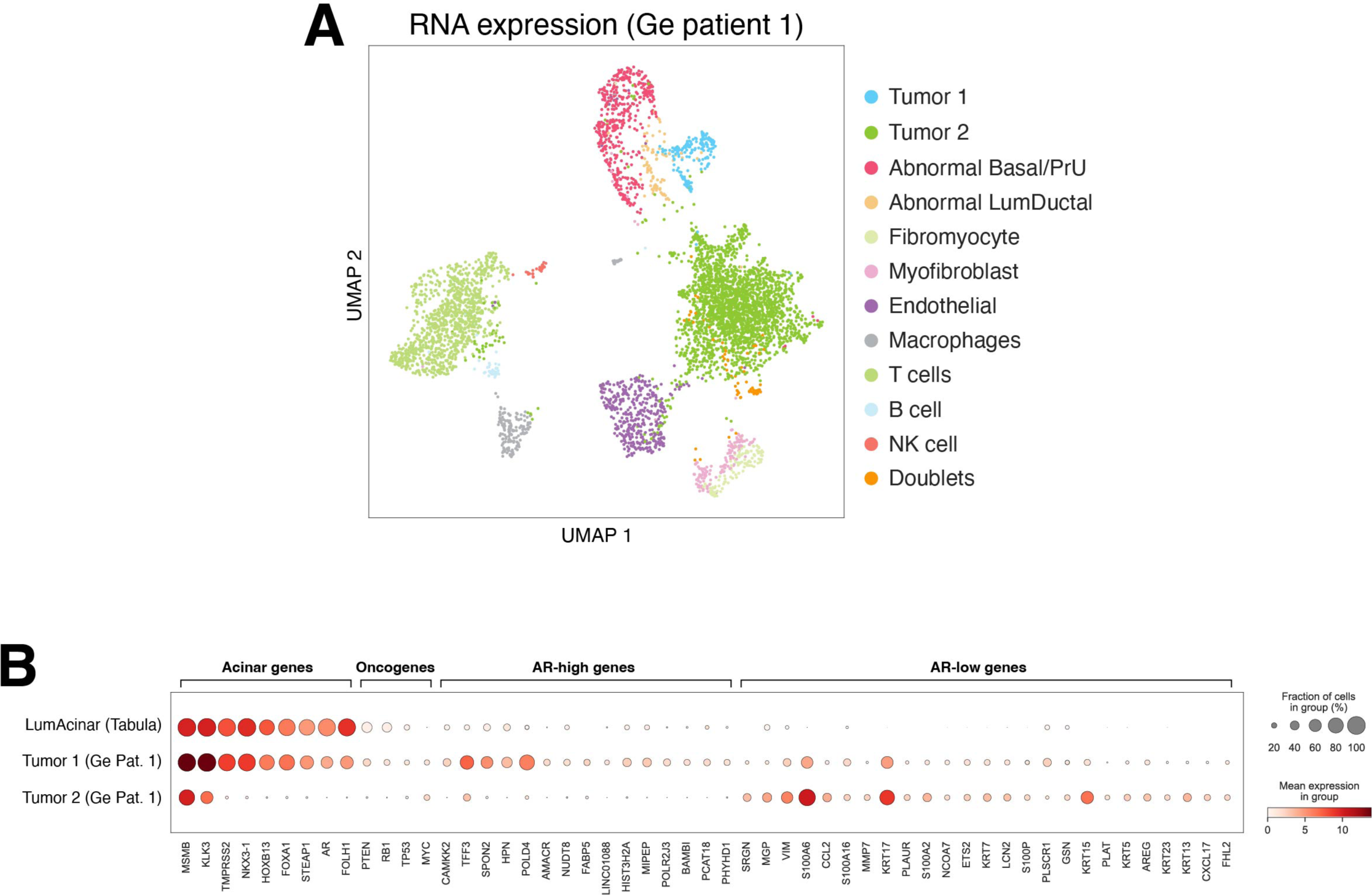
Full InferCNV analysis of the patient 1 tumor from the Ge cohort. **(A)** UMAP plots of gene expression for cell populations from patient 1 ***(Ge et al., 2022)***, showing two distinct tumor populations and multiple abnormal epithelial cell types. **(B)** Dot plot of the normal LumAcinar cells (Tabula), and Patient 1 LumAcinar Tumor 1 and Tumor 2 cells, showing that Patient 1 had both AR-high and AR-low tumor cells concurrently.

**Figure 6—source data 1.** Human prostate samples and corresponding clinical data.

| Sample ID | Diagnosis | Gleason Grade | Gleason Score (highest) | Stage | Age | Race | Ethnicity | Treatments | PSA at sample acquisition | Time of PSA (months before acquisition) | PSA peak |
| --- | --- | --- | --- | --- | --- | --- | --- | --- | --- | --- | --- |
| LHu1 | Adenocarcinoma | GG3 | 4+3 and 3+3 | pT2N0 | 65 | White | Non-Hispanic | None | 9 | 6 | 9 |
| LHu2 | Adenocarcinoma | GG1 | 3+3 | pT2N0 | 57 | Combination Not Described | Hispanic | None | 7 | 2 | 7 |
| LHu3 | Adenocarcinoma (metastatic) | GG5 | 4+5 | ypT3N1 | 67 | White | Non-Hispanic | Degarelix | 36 | 0 | 94 |
| LHu4 | Adenocarcinoma | GG2 | 3+4 | pT3N0 | 64 | Combination Not Described | Hispanic | None | 27 | 2 | 27 |
| LHu5 | Adenocarcinoma | GG4 | 4+4 | pT2N0 | 69 | Combination Not Described | Non-Hispanic | Tamsulosin | 12 | 19 | 12 |

